# Dually targeted proteins regulate proximity between peroxisomes and partner organelles

**DOI:** 10.1101/2022.07.29.501968

**Authors:** Thorsten Stehlik, Elena Bittner, Jason Lam, Lazar Dimitrov, Isabelle Schöck, Jannik Harberding, Nikola Heymons, Maya Schuldiner, Einat Zalckvar, Michael Bölker, Randy Schekman, Johannes Freitag

## Abstract

Peroxisomes play a central role in fatty acid metabolism. To correctly target to peroxisomes, proteins require specialized targeting signals. One mystery in the field is sorting of proteins that carry both a targeting signal for peroxisomes as well as for other organelles such as mitochondria or the endoplasmic reticulum (ER). Exploring several of these dually localized proteins in *Saccharomyces cerevisiae,* we observed that they can act as dynamic tethers bridging organelles together through an affinity for organelle-destined targeting factors. We show that this mode of tethering involves the peroxisome import machinery, the ER– mitochondria encounter structure (ERMES) in the case of mitochondria and the GET complex in the case of the ER. Depletion of each of the targeting factors resulted in the accumulation of smaller peroxisomes. We propose that dual targeting of proteins occurs at contact sites and that protein import *per se* contributes to the maintenance of these membrane proximities. This introduces a previously unexplored concept of how targeting of dual affinity proteins can support organelle attachment, growth and communication.

## Introduction

Metabolic pathways in eukaryotic cells are often compartmentalized inside membrane-enclosed organelles such as mitochondria and peroxisomes [1]. Localized proteins synthesized in the cytosol must target and translocate into their organelles of residence correctly, efficiently and in a regulated manner [2–4]. The majority of mitochondrial proteins are imported from the cytosol in a conformationally flexible state via evolutionary conserved protein complexes called translocase of the outer membrane (TOM) and translocase of the inner mitochondrial membrane (TIM). Many proteins destined for the mitochondrial matrix or inner membrane contain N-terminal targeting signals that are cleaved by proteases upon import (Wiedemann & Pfanner, 2017).

Protein translocation into the matrix of peroxisomes requires distinct targeting signals termed peroxisome targeting signal type 1 and 2 (PTS1 and PTS2, respectively). PTS1 motifs are recognized by the cytosolic targeting factor Pex5 and translocated via interaction of Pex5 with Pex13/14 on the peroxisomal membrane allowing the translocation of folded and even oligomerized protein complexes in a yet unresolved mechanism [4,6,7]. Peroxisomes can multiply by growth and division, but unlike mitochondria can also form *de novo* in the absence of mature peroxisomes from the endoplasmic reticulum (ER) and possibly also involving mitochondria [4,8,9].

For many years proteins were studied that either have a C-terminal PTS1 or an N-terminal mitochondrial targeting signal (MTS). Recently, it was shown that the protein phosphatase Ptc5 of the yeast *S. cerevisiae* (from hereon called simply yeast) contains both targeting signals and exhibits a dual localization to peroxisomes and mitochondria [10]. Ptc5 is processed by the inner membrane peptidase (IMP) complex inside mitochondria and reaches the peroxisomal matrix via mitochondrial transit [10, 11]. Additional proteins were identified that contain competing mitochondrial and peroxisomal targeting signals. Overexpression of several enhanced the number of peroxisomes in proximity to mitochondria [10]. Also data from mammals point to a similar mode of tethering [12].

Organelle tethering has been a subject of intense research in the last years as it has become clear that organelles do not work in isolation but rather form areas of close apposition, contact sites, that enable the transfer of molecules [13]. For several years now tethering molecules that sustain such contacts have been discovered and, until now, have been shown to rely on components that either span the membrane or tightly bind to it [14]. One such contact site is formed between peroxisomes and mitochondria and has been dubbed the PerMit [15, 16]. Genetic data and protein interaction experiments suggests that the peroxisomal membrane protein Pex11 is involved in forming this contact site [16, 17]. Other tethers for peroxisomes have been identified [18–20] e.g. Inp1, important for partitioning of peroxisomes upon cell division in yeasts [21–23]. In mammalian cells, proximity of peroxisomes and the ER is achieved through a pair of membrane-associated proteins assisting lipid transfer [24, 25]. Since deleting any single tether does not result in loss of the contact site it is clear that multiple tethering mechanisms underlie the formation of each contact but, for the most part, these have not been worked out.

Here, we report on dynamic tethers that are formed by proteins with affinity to two targeting/translocation machineries. We propose a model suggesting that sorting of cargo can not only occur at organelle contact sites but also can contribute to their stabilization. This concept establishes dual targeting as a way to dial the exact amount of proximity required between cellular compartments.

## Results

### Proteins that contain both an MTS and a PTS1 increase proximity of mitochondria and peroxisomes upon overexpression

A diverse group of proteins contain dual targeting information for peroxisomes and mitochondria [10, 26] (Supplementary table 1). Owing to their characteristic domain structure (Fig. 1A), with targeting signals at both termini, such proteins may be able to simultaneously associate with both organelles and become distributed at their interface. Hence, their sorting may either require, contribute or both to organelle contact.

**Figure 1.**
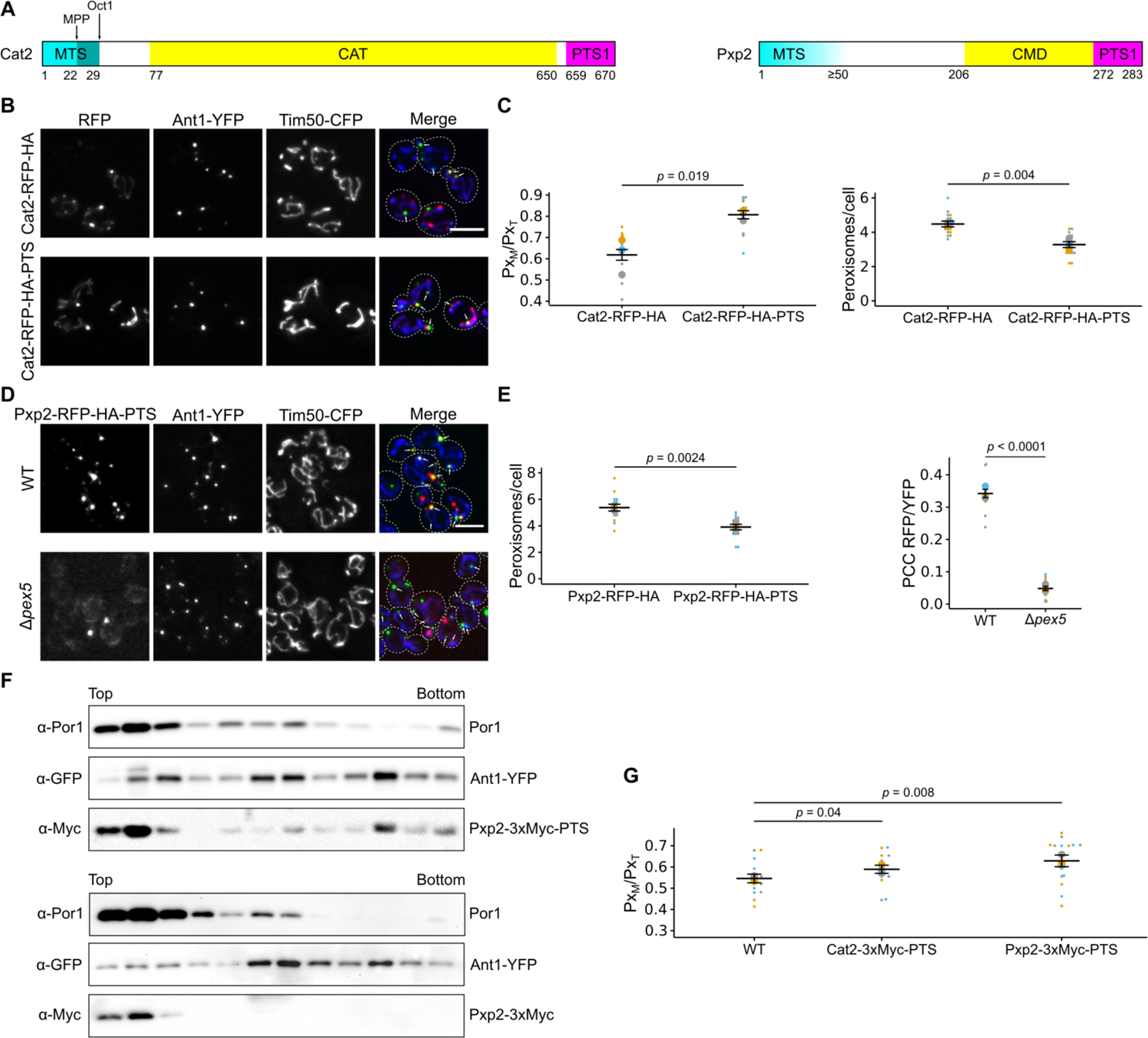
Cat2 and Pxp2 localize to mitochondria and peroxisomes and induce interorganellar contacts. **(A)** Scheme showing the domain architecture of Cat2 and Pxp2. Cat2 contains an N-terminal mitochondrial targeting signal (MTS) which is likely processed sequentially by two different mitochondrial proteases (MPP, Oct1) (Morgenstern et al., 2017) and contains an acetyl-carnitine transferase (CAT) domain and a PTS1 (Elgersma et al., 1995). Pxp2 contains a N-terminal MTS, a putative carboxymuconolactone decarboxylase domain (CMD) and a PTS1 (Nötzel et al., 2016, Stehlik et al., 2020). Prediction of protein domains was performed with the HMMER web server (https://www.ebi.ac.uk/Tools/hmmer/) **(B)** Fluorescence microscopic images of wildtype yeast cells expressing either Cat2-RFP-HA or Cat2-RFP-HA-PTS (red) together with the peroxisomal membrane protein Ant1-YFP (green) and the mitochondrial inner membrane protein Tim50-CFP (blue). White arrows denote peroxisomes overlapping mitochondria **(C)** Quantification of the fraction of peroxisomes in contact with mitochondria (Px_M_) relative to the total peroxisome number (Px_T_) (left) and quantification of the number of peroxisomes per cell (right) of cells shown in (B). **(D)** Fluorescence microscopic pictures of wildtype (WT) and Δ*pex5* cells co-expressing Pxp2-RFP-HA-PTS together with Ant1-YFP and Tim50-CFP (left). White arrows denote peroxisomes overlapping mitochondria. Scale bar represents 5 µm. **(E)** Quantification of the number of peroxisomes per cell (left) of cells and correlation between the Pxp2-RFP-HA-PTS signal and Ant1-YFP signal (right) of cells shown in (D). PCC refers to Pearson’s correlation coefficient. **(F)** Subcellular localization of endogenously tagged Pxp2-3xMyc-PTS and Pxp2-3xMyc was determined using density gradient centrifugation. Twelve fractions, collected from the top of the gradient, were analyzed by SDS–PAGE and immunoblot. Ant1-YFP is a peroxisomal membrane protein and Por1 is localized in the mitochondrial outer membrane. **(G)** Quantification of the fraction of peroxisomes contacting mitochondria (Px_M_) in relation to the total peroxisome count (Px_T_) of wildtype (WT) cells and cells overexpressing the indicated fusion proteins. Scale bars represent 5µm. Quantifications are based on *n* = 3 experiments. Each color represents one experiment. Error bars represent SEM. *P*-values were calculated using a two-sided unpaired Student’s *t*-test.

Overexpression of several such candidates (Cat2, Dpi8, Mss2, Pxp2, Tes1) fused to a Red Fluorescent Protein (RFP) on their C-terminus, in a manner preserving the original PTS1, elevated the number of peroxisomes in proximity to mitochondria (Figs. 1A - 1C and S1; Supplementary table 1) [10]. Targeting to peroxisomes was dependent on the presence of the targeting factor Pex5 (Figs.1D and S1). Overexpression of Pxp2-RFP-PTS and Cat2-RFP-PTS reduced the total peroxisome number (Fig. 1C and 1E). Both results may be explained by a potential tethering function.

We considered the possibility that enhanced proximity between the organelles may have resulted from the use of a bulky RFP tag which is known in other cases to affect import into organelles [10]. To test this, we internally Myc-tagged Cat2, Cta1, Pxp2 and Tes1 at their endogenous loci maintaining their original N- and C-termini. Dual targeting was confirmed by evaluation of compartments resolved on buoyant density gradients (Figs. 1F and S2).

We then utilized overexpressed internally Myc-tagged versions of two candidate proteins integrated into the *LEU2* locus: Cat2 and Pxp2. Even without the RFP tag, Pxp2-Myc-PTS and Cat2-Myc-PTS both increased the association of peroxisomes and mitochondria (Fig. 1G) with Pxp2 giving a stronger effect. A difference between both proteins was also reflected by their subcellular distribution. Whereas Cat2-RFP-PTS was found to localize distributed over mitochondria and in peroxisomes, Pxp2-RFP-PTS occurred in foci at mitochondria frequently proximal to a peroxisome but also inside peroxisomes with no contact to mitochondria (Fig. 1B and 1D). Pxp2-RFP-PTS became more evenly distributed in mitochondria upon deletion of *PEX5* (Fig. 1D). Thus, overexpression of proteins that have a mitochondrial and a peroxisomal targeting signal can enhance the formation of contacts between both organelles.

### Loss of the matrix protein targeting machinery reduces the number of contacts between peroxisomes and mitochondria

We reasoned that if tethering is caused by dual targeting then we should be able to affect it by eliminating the targeting factor Pex5. We therefore quantified the number of peroxisomes associated with mitochondria in control and Δ*pex5* cells. A significant reduction of peroxisomes proximal to mitochondria was observed in the absence of Pex5 (Fig. 2A and 2B) indicating a role of the peroxisomal matrix targeting machinery for the generation of contacts – something unexpected if only membrane proteins were required for tethering as is usually assumed. Indeed, we could block enhanced tethering triggered by overexpression of Pxp2-RFP-PTS1 by deleting *PEX5* (Fig. 2C). Furthermore, we quantified peroxisome number and detected an increase of smaller appearing peroxisomes upon deletion of *PEX5* (Fig. 2D). Increased movement of peroxisomes was detected in Δ*pex5* cells (Movie 1), a phenotype previously described for other mutants with reduced organelle tethering [e.g. ref 24]. These data show that overexpression of dual affinity PTS1 proteins and depletion of Pex5 have a reciprocal effect on peroxisome number and appearance.

**Fig. 2.**
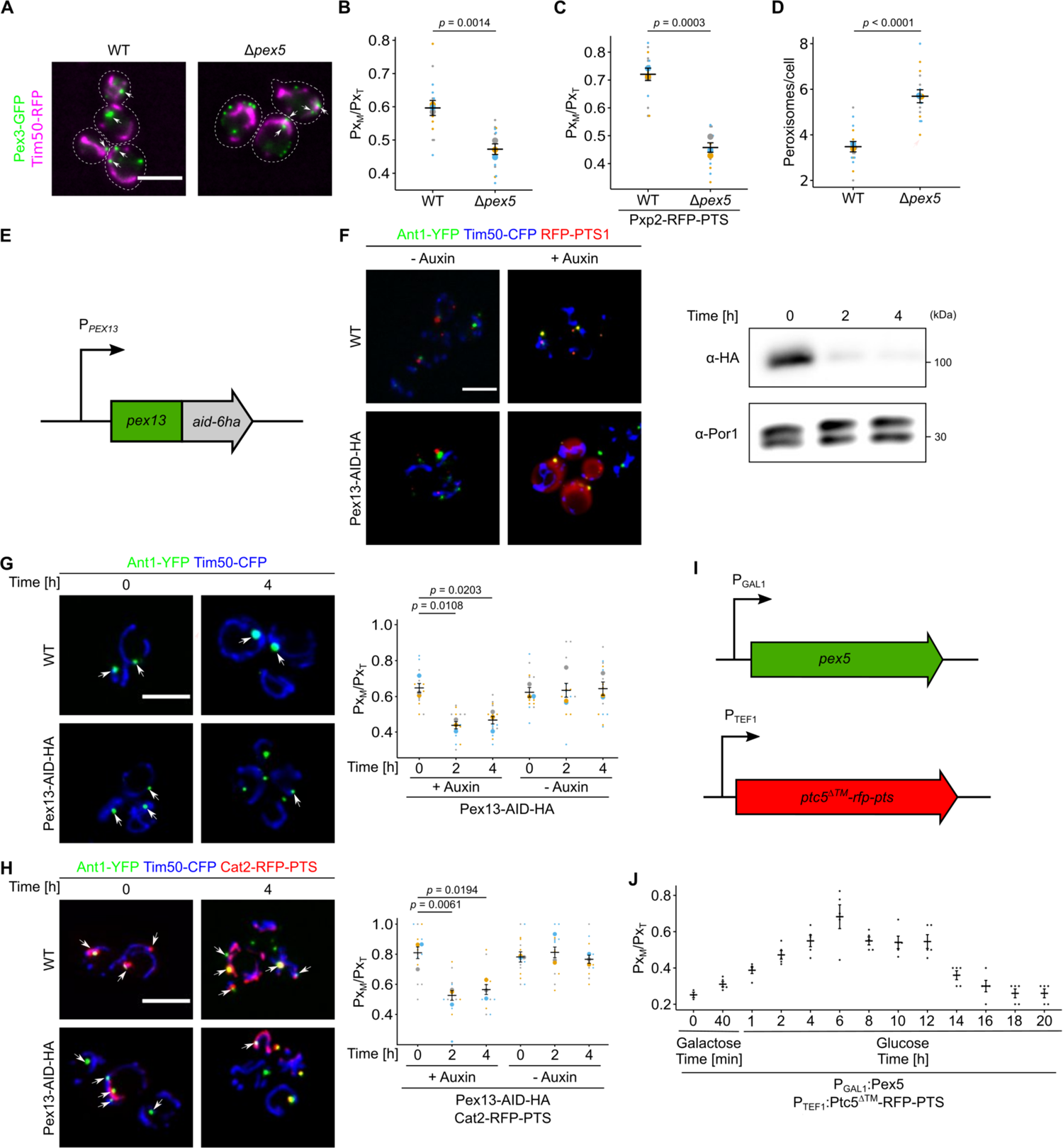
Depletion of components of the peroxisomal import machinery reduces PerMit contacts. **(A)** Fluorescence microscopic picture from WT and Δ*pex5* cells expressing endogenously tagged Pex3-GFP and Tim50-RFP. White arrows denote peroxisomal signal overlapping with mitochondrial signal. **(B)** Quantification of the fraction of peroxisomes contacting mitochondria (Px_M_) in relation to the total peroxisome count (Px_T_) of wildtype (WT) cells and Δ*pex5* cells from A. The number of peroxisomes per cell was quantified in the indicated strains (right) **(C)** Quantification of the fraction of peroxisomes contacting mitochondria (Px_M_) in relation to the total peroxisome count (Px_T_) of wildtype (WT) cells and Δ*pex5* cells expressing Pxp2-RFP-PTS, Ant1-YFP and Tim50-CFP. **(D)** The number of peroxisomes per cell was quantified in the indicated strains expressing Pex3-GFP. **(E)** Scheme of the genetic modifications used for auxin-dependent depletion of Pex13 in (F), (G) and (H). The endogenous *PEX13* locus was genetically engineered to encode a translational fusion of Pex13 with a C-terminal auxin-inducible degron (AID) and six hemagglutinin (HA) tags. Pex13 degradation is mediated by the F-box protein AFB2 from A. *thaliana*, which was expressed from the *ADH1* promotor **(F)** Fluorescence microscopic images of indicated strains expressing the peroxisomal membrane protein Ant1-YFP (green), the mitochondrial inner membrane protein Tim50-CFP (blue) and RFP-PTS (red) after a 4 h incubation period in the absence (-Auxin) or presence (+Auxin) of 2 mM indole-3-acetic acid (left). Auxin-dependent depletion of Pex13-AID-HA at indicated time points was analyzed by SDS-PAGE and immunoblot (right). Por1 served as a loading control. **(G)** Subcellular localization of Ant1-YFP and Tim50-CFP of indicated strains was analyzed in the presence of 2 mM indole-3-acetic acid at indicated time points (left). Arrows indicate peroxisomes in close proximity to mitochondria. The fraction of peroxisomes in contact with mitochondria (Px_M_) relative to the total peroxisome count (Px_T_) of the indicated strain was quantified at the indicated time points and growth conditions (right). **(H)** Identical to (G), except that the cells also expressed Cat2-RFP-PTS1 to increase PerMit contacts. **(I)** Schematic illustration of a galactose-inducible conditional *pex5* mutant overexpressing Ptc5^ΔTM^-RFP-PTS to generate additional PerMit contacts. **(J)** Quantification of the fraction of peroxisomes in contact with mitochondria (Px_M_) relative to the total peroxisome count (Px_T_) of the strain described in (I) at the indicated time points and growth conditions from images derived from time lapse microscopy. Scale bars represent 5 µm. Quantifications are based on *n* = 3 experiments. Each color represents one experiment. Error bars represent SEM. *P*-values were calculated using a two-sided unpaired Student’s *t*-test.

To confirm our findings and reduce the possibility that the observed phenotype resulted from secondary, downstream effects we took an approach for rapid depletion of the Pex5 receptor Pex13 [7], using an auxin-inducible degron (AID) tag [27, 28] (Fig. 2E). Depletion of Pex13 resulted in reduced import of RFP-PTS1 (Fig. 2F). Association of peroxisomes and mitochondria was quantified in a strain co-expressing Cat2-RFP-PTS as an additional tether relative to a control strain (Fig. 2G and 2H, Fig. S3).

Both strains started with a different degree of organelle association but depletion of Pex13-AID-HA by auxin addition reduced the number of peroxisomes proximal to mitochondria to a similar basal level supporting a role of matrix protein targeting in driving organelle association. To corroborate these results we created a strain with conditional *PEX5* expression by putting it under control of the *GAL* promoter (Fig. 2I). Immunoblot and fluorescence microscopy of RFP-PTS1 expressing cells confirmed that Pex5 levels correlated with PTS1 import (Fig. S3). We then overexpressed a Ptc5ΔTMD-RFP-PTS, which is a dual targeted derivative of Ptc5 lacking the transmembrane domain [10]. Automated time lapse imaging revealed that induction of Pex5 elevated the number of peroxisomes close to mitochondria, whereas its reduction had the opposite effect (Fig. 2J). Since single planes were imaged, lower basal overlap of mitochondria and peroxisomes were observed in the absence of Pex5 (compare to Fig. 2B). Together, these data suggest a function of peroxisomal matrix protein import for formation of PerMit contacts.

### Identification of factors that regulate dual targeting of Ptc5

To identify additional factors involved in dual targeting and therefore potentially also tethering, we conducted a high content genetic screen focused on Ptc5 sorting from mitochondria to peroxisomes (Fig. 3A). Peroxisomal import of Ptc5-RFP-PTS depends on mitochondrial processing and subsequent Pex5 dependent targeting [10]. This transit mechanism may require direct contact between the organelles. Crossings of a wildtype strain expressing Ptc5-RFP-PTS and Pex3-GFP as a peroxisomal marker with arrayed libraries containing yeast deletion strains and hypomorphic mutants of essential genes were performed and sporulated to select for haploid cells containing both fluorescent proteins together with a genetic perturbation (Fig. 3A) [29, 30]. Inspection of haploid progeny by automated microscopy uncovered various genes affecting peroxisomal targeting of Ptc5-RFP-PTS (Fig. 3B and 3C; Supplementary table 2). These include many expected genes essential for peroxisome biogenesis but also distinct genes e.g. *SOM1*. Som1 was shown previously to be part of the IMP complex [31], which we could confirm (Fig. 3D). Moreover, we identified Pex34 as a factor for correct sorting of Ptc5-RFP-PTS to peroxisomes (Fig. 3C). Pex34 was identified as a potential PerMit tether [16].

**Figure 3.**
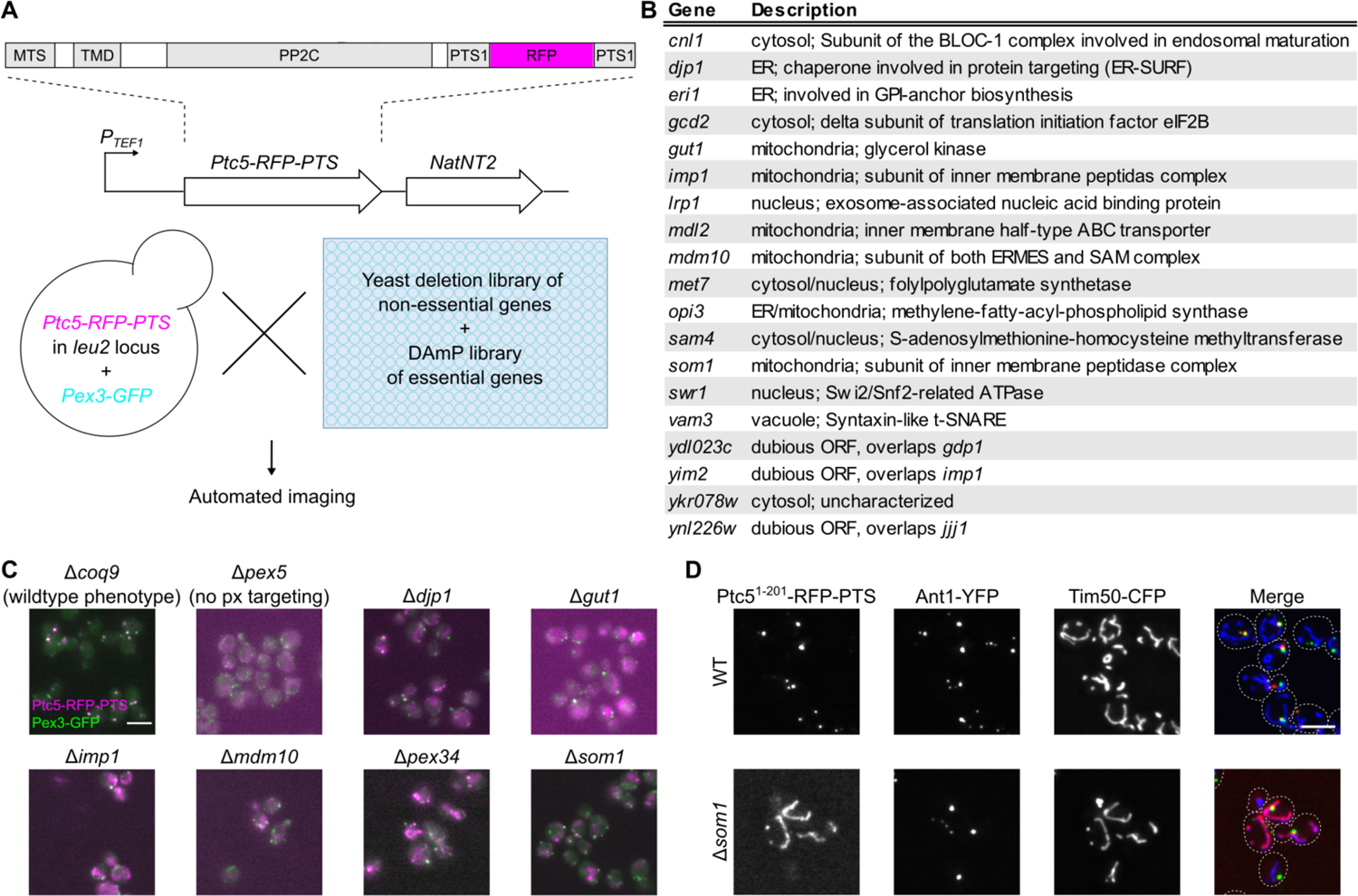
Identification of factors involved in targeting of Ptc5 to peroxisomes by a high content genetic screen. **(A)** Schematic illustration of a high-content microscopic screen aimed to uncover factors involved in sorting of the protein phosphatase Ptc5. A query strain co-expressing Ptc5-RFP-PTS and Pex3-GFP was crossed into indicated yeast libraries using synthetic genetic array technology. Haploid progeny expressing both fusion proteins in each mutant background were analyzed with automated fluorescence microscopy. MTS: mitochondrial targeting signal, TMD: transmembrane domain, PP2C: protein phosphatase type 2C, PTS1: peroxisomal targeting signal type 1. **(B)** Table showing a subset of mutants affecting Ptc5 sorting. The entire list of mutations is depicted in Supplementary table 2. **(C)** Subcellular localization of Ptc5-RFP-PTS (magenta) and Pex3-GFP (green) was analyzed in indicated strains using automated fluorescence microscopy. Δ*coq9* cells were used as a control as these show Ptc5-RFP-PTS1 localization in peroxisomes, but are affected in mitochondrial metabolism (Stehlik et al., 2020). Δ*pex5* mutants show no peroxisomal targeting of the reporter Ptc5-RFP-PTS1 but only mitochondrial signal. **(D)** Fluorescence microscopic images of cells co-expressing the truncated Ptc5 allele Ptc5^1-201^-RFP-PTS (red) (Stehlik et al., 2020) together with Ant1-YFP (green) and Tim50-CFP (blue) in indicated strains. Scale bars represents 5 µm. This result strengthen our hypothesis that the proximity of mitochondria and peroxisomes is crucial for peroxisomal targeting of Ptc5-RFP-PTS via mitochondrial transit and that our assay is likely to report on tethering.

### The ERMES complex regulates dual targeting between mitochondria and peroxisomes

An interesting gene target from the screen was Mdm10 (a critical factor for connecting mitochondria to the ER as part of the ER-mitochondria encounter structure (ERMES)) (Figs. 3 and 4A). The ERMES complex consists of the ER-anchored protein Mmm1 and three mitochondrial proteins (Mdm10, Mdm12 and Mdm34) [32]. The integral membrane protein Mdm10 has an additional function in assisting import of mitochondrial ß-barrel proteins [33]. It was therefore unclear whether the effect of Mdm10 was through the ERMES or by affecting mitochondrial protein import.

**Figure 4.**
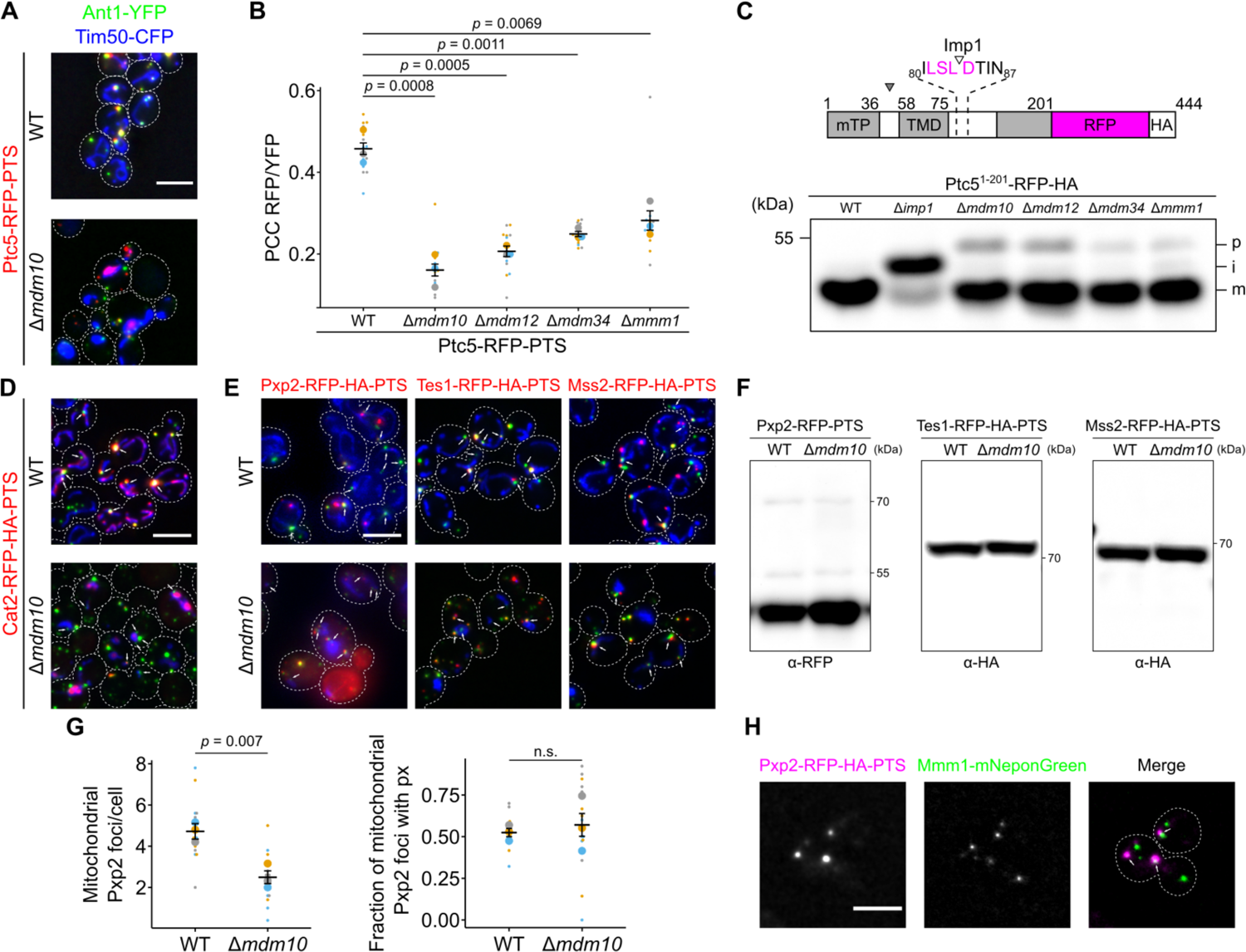
ERMES complex regulates import of proteins into mitochondria and peroxisomes. **(A)** Subcellular localization of Ptc5-RFP-PTS (red), the peroxisomal membrane protein Ant1-YFP (green) and the mitochondrial inner membrane protein Tim50-CFP (blue) in indicated strain was analyzed with fluorescence microscopy. **(B)** Correlation between Ptc5-RFP-PTS signal and Ant1-YFP signal was quantified in indicated strains. PCC refers to Pearson’s correlation coefficient. **(C)** The truncated variant Ptc5^1-201^-RFP lacking a PTS1 was expressed in indicated strains. Cleavage sites for MPP (filled arrow) and the IMP complex (blank arrow) are indicated in the scheme. Whole cell lysates were analyzed by SDS-PAGE and immunoblot. p: premature isoform, i: intermediate isoform, m: mature isoform. Concentrations of protein extracts were adapted to each other to focus on processing. **(D)** Cat2-RFP-PTS (red) was co-expressed with Ant1-YFP (green) and Tim50-CFP (blue) in wildtype (WT) or Δ*mdm10* cells. Subcellular localization was determined with fluorescence microscopy. White arrows denote peroxisomes overlapping mitochondria. **(E)** Fluorescence microscopic pictures of indicated strains expressing respective RFP fusion proteins (red) together with Ant1-YFP (green) and Tim50-CFP (blue). White arrows denote peroxisomes overlapping mitochondria. **(F)** Whole cell lysates of strains expressing the indicated fusion proteins were analyzed by SDS-PAGE and immunoblot. Concentrations of protein extracts were adapted to each other to focus on processing. **(G)** The number of Pxp2-positive foci per cell at mitochondria was quantified in indicated strains (left). Quantification of the fraction of mitochondrial Pxp2 foci overlapping Ant1-YFP (right). **(H)** Fluorescence microscopic picture of a strain expressing Pxp2-RFP-PTS (magenta) together with Mmm1-mNeon (green). White arrows indicate Pxp2-RFP-PTS foci overlapping Mmm1-NeonGreen foci. Scale bars represent 5 µm. Quantifications are based on *n* = 3 experiments. Each color represents one experiment. Error bars represent SEM. *P*-values were calculated using a two-sided unpaired Student’s *t*-test.

To address whether the observed phenotype is specific to Mdm10 or shared by other ERMES complex members, we repeated our assay in freshly made strains. In all ERMES mutants targeting of Ptc5-RFP-PTS to peroxisomes was significantly reduced (Figs. 4B and S4A) with *Δmdm10* displaying the strongest phenotype.

To test the stage at which ERMES mutants affected the dually targeted proteins, we first analyzed the initial steps of mitochondrial protein import. All mutants influenced mitochondrial import and preprotein processing as determined by testing a truncated variant of Ptc5-RFP (Fig. 4C). Moreover, we observed an import defect for several *bona fide* mitochondrial proteins in Δ*mdm10* cells (Fig. S4B) demonstrating that loss of ERMES complex alters translocation of mitochondrial proteins in a more general way.

To follow up on these results, we determined the localization of other proteins that contain competing targeting signals in Δ*mdm10* cells. Cat2-RFP-PTS behaved similar to Ptc5-RFP-PTS and was retained in mitochondria while targeting to peroxisomes was reduced (Fig. 4D). In contrast, the localization of Pxp2-RFP-PTS, Tes1-RFP-PTS and Mss2-RFP-PTS displayed the opposite pattern of localization in Mdm10 depleted cells with reduced mitochondrial targeting but normal peroxisome targeting (Fig. 4E). Proteolytic processing defects could not be observed for any of these proteins suggesting a noncanonical targeting signal for mitochondria (Fig. 4F). Quantification showed a reduction of mitochondrial foci containing Pxp2-RFP-PTS (Fig. 4G). Proteins containing a classical mitochondrial presequence such as Ptc5 may therefore require ERMES-mediated proximity to translocate to peroxisomes, whereas proteins without a canonical signal do not efficiently target mitochondria in the absence of ERMES. Given these results, we suggest that previous literature showing that overexpression of Mdm10 and Gem1 enhanced PerMit contacts [16] exposed their ability to provide excess of dual affinity proteins to mitochondria. ERMES components could be directly involved in mitochondrial targeting as we found that Pxp2-RFP-PTS foci regularly (38% +/- 5%) overlapped with Mmm1-mNeonGreen foci (Fig. 4H; Supplementary table 3). An indirect function of ERMES is likewise possible and a potential saturation of the import machinery on the mitochondrial surface provokes the altered localization of dual affinity proteins such as Pxp2, Mss2 and Tes1.

### ERMES controls proximity of peroxisomes and mitochondria

Quantification of peroxisomes in all ERMES mutants showed that they contained a higher number of small peroxisomes in a manner similar to a *Δpex5* strain (Fig. 2) and that import of RFP-PTS1 was reduced (Fig. 5A). Moreover, the fraction of peroxisomes in contact to mitochondria declined in the ERMES mutants (Fig. 5B and 5C). These phenotypes often reverted and, for example, a Δ*mdm12* deletion strain with wildtype like growth and regular mitochondrial morphology retained a higher number of peroxisomes (Fig. S5).

**Figure 5.**
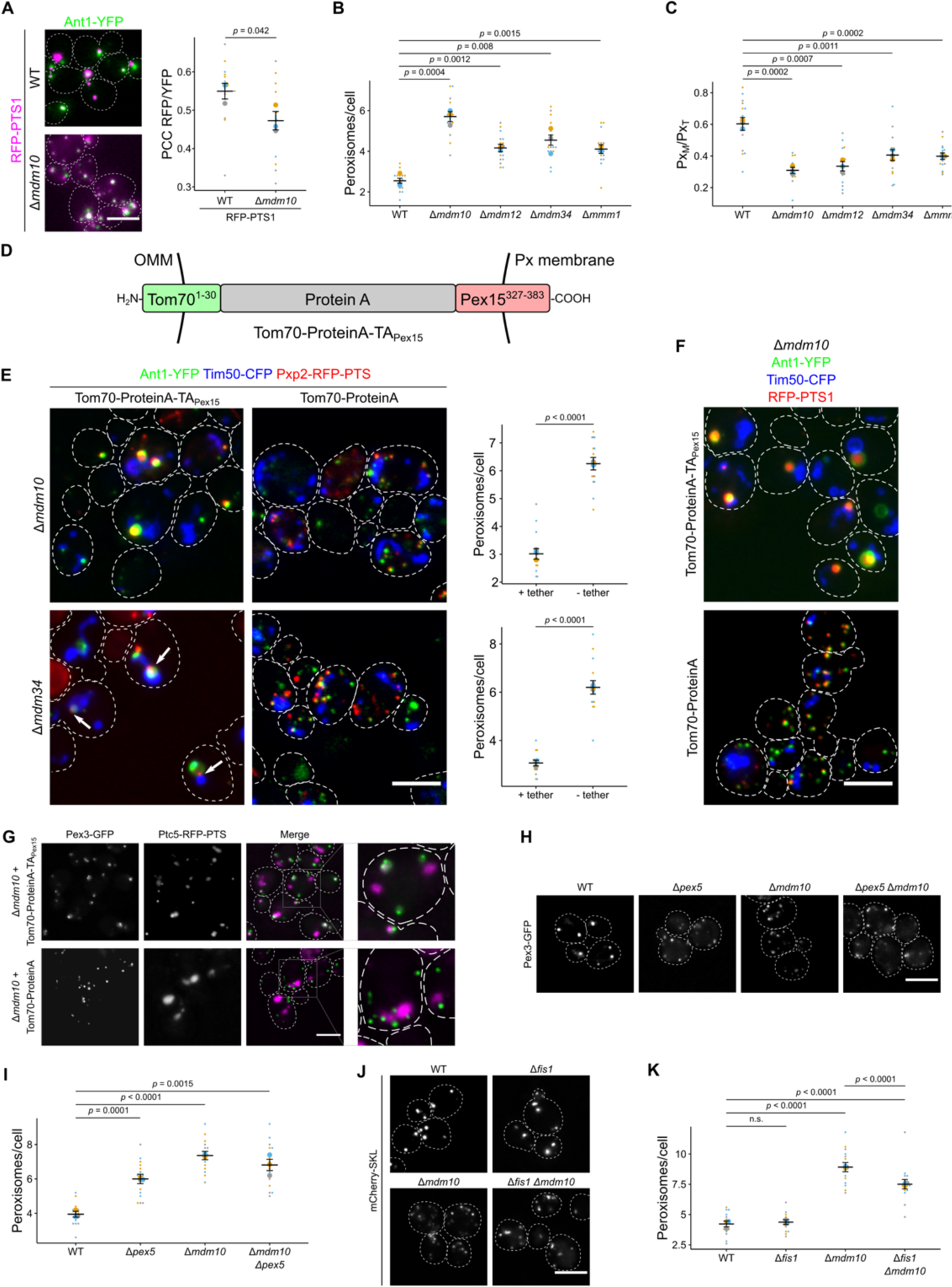
Peroxisome function and proximity to mitochondria is affected in ERMES mutants. **(A)** Fluorescence microscopic images of wildtype (WT) and Δ*mdm10* cells co-expressing the peroxisomal marker RFP-PTS1 (magenta) and the peroxisomal membrane protein Ant1-YFP (green) (left). Correlation of the RFP-PTS1 signal and the Ant1-YFP signal was quantified using Pearson’s correlation coefficient (PCC; right). **(B)** The number of peroxisomes per cell was quantified in indicated strains. **(C)** Quantification of the ratio of peroxisomes in contact with mitochondria (Px_M_) to the total peroxisome count (Px_T_) **(D)** Scheme of the synthetic PerMit tether. OMM: outer mitochondrial membrane, Px: peroxisome, TA: tail-anchor **(E)** A synthetic tether for peroxisomes and mitochondria (Tom70-ProteinA-TA_Pex15_) can suppress accumulation of small peroxisomes. Fluorescence microscopic images of strains deleted for *MDM10* (upper panel) or *MDM34* (lower panel) co-expressing Pxp2-RFP-PTS (red) and the marker proteins Ant1-YFP (green) and Tim50-YFP (blue) in presence of the tether Tom70-ProteinA-TA_Pex15_ or a control protein (left). Quantification of the number of peroxisomes per cell of the indicated strains (right). Arrows indicate Pxp2-RFP-PTS foci located at junctions between mitochondria and peroxisomes in Δ*mdm34* cells. **(F)** Images of Δ*mdm34* cells co-expressing RFP-PTS1 (red) together with Ant1-YFP (green), Tim50-CFP (blue) and Tom70-ProteinA-TA_Pex15_ or a control protein. **(G)** Representative pictures of Δ*mdm10* cells expressing Ptc5-RFP-PTS (magenta) integrated into the *LEU2* locus and Pex3-GFP (green) together with the tether Tom70-ProteinA-TA_Pex15_ or a control protein. Magnifications show enlarged peroxisomes containing Ptc5-RFP-PTS upon expression of the synthetic tether. Fluorescence microscopic pictures of cells expressing Pex3-GFP **(H)** or mcherry-SKL **(J).** The number of peroxisomes expressing Pex3-GFP **(I)** or mCherry-SKL **(K)** was quantified. Scale bars represent 5µm. Quantifications are based on *n* = 3 experiments. Each color represents one experiment. Error bars indicate SEM. *P*-value was calculated using a two-sided unpaired Student’s *t*-test.

Indeed, phenotypic suppression of ERMES mutants by second site mutations is regularly observed [34]. However, the effect of ERMES loss on peroxisome number, proximity to mitochondria and matrix targeting speaks for a specific function of ERMES proteins on peroxisome formation independent of mitochondrial dysfunction as was also suggested elsewhere [17, 35].

The above led us to consider that ERMES may affect the proximity of mitochondria and peroxisomes – directly or indirectly. Since we had noticed that overexpression of dual-affinity proteins resulted in increased proximity and a reduced number of peroxisomes (Fig. 1), we tested whether a synthetic tether – designed to connect peroxisomes and mitochondria (Fig. 5D) also affected peroxisomes in a manner that rescued loss of ERMES components. We found that in both Δ*mdm10* and Δ*mdm34* cells expression of a synthetic tether decreased peroxisome number (Figs. 5E and S6). In addition, expression of this synthetic tether activated substantial expansion of peroxisomes (Fig. 5E and 5F).

Upon deletion of *MDM34* (and less so in *Δmdm10*) we were able to visualize Pxp2-RFP-PTS, but not the soluble peroxisomal marker protein RFP-PTS (Figs. 5F and S6) accumulated at junctions between mitochondria and the enlarged peroxisomes (Figs. 5E and S6). This observation supports the function of Pxp2-RFP-PTS as a dual affinity tether distributed at sites of organelle contact. Furthermore, expression of the synthetic tether partially restored peroxisomal targeting of Ptc5-RFP-PTS in Δ*mdm10* cells (Fig. 5G). These findings strengthen a proposed role of ERMES as a regulator of the interaction of mitochondria and peroxisomes. Deletion of *PEX5* and deletion of *MDM10* both resulted in a higher number of small peroxisomes, with Δ*mdm10* having the stronger effect (Fig. 5H and 5I). The phenotype of the double mutant was more reminiscent to Δ*mdm10* single mutants in line with a potential connection of both factors (Fig. 5H and 5I). The stronger phenotype of the Δ*mdm10* deletion strain points to additional functions of ERMES for peroxisome formation beside its role in regulating trafficking of dual affinity PTS1 proteins. These may include stabilization of contacts via interaction with membrane proteins such as Pex11 as previously suggested [17] or enhanced activatity of fission proteins such as the dually localized tail-anchored (TA) fission protein Fis1 on peroxisomes [36] in cells lacking ERMES. Indeed, additional deletion of *FIS1* partially suppressed peroxisome accumulation in Δ*mdm10* cells (Fig. 5J and K).

### The TA-protein Pex15 contributes to ER-peroxisome proximities

In analogy to dual targeting of Cat2-RFP-PTS, Ptc5-RFP-PTS and Pxp2-RFP-PTS1, sorting of dually targeted membrane proteins could occur at organelle contact sites and may contribute to their formation or stabilization. Many peroxisomal membrane proteins localize to the ER or mitochondria in peroxisome deficient strains lacking the chaperone Pex19 or the membrane protein Pex3 [37, 38]. One such protein is the TA protein Pex15 (Pex26 in mammals) which can be sorted via the ER as well as directly imported into peroxisomes via Pex19 [39–44]. Pex15 can be targeted and translocated into the ER via the GET complex [39]. We therefore expressed the reporter protein ProteinA (PA)-GFP-Pex15g containing a C*-*terminal glycosylation tag (g) under control of the *MET25* promoter in the presence of 200 µM methionine. The chimeric fusion protein localized predominantly in peroxisomes of wildtype cells or upon deletion of *get3* and largely in the ER in *Δpex19* cells (Fig. 6A and 6B). This behavior is similar to several dual affinity PTS1 cargo in Δ*pex5* and Δ*mdm10* mutants, respectively (compare to Figs. 1, S1 and 4D; Stehlik et al., 2020).

**Figure 6.**
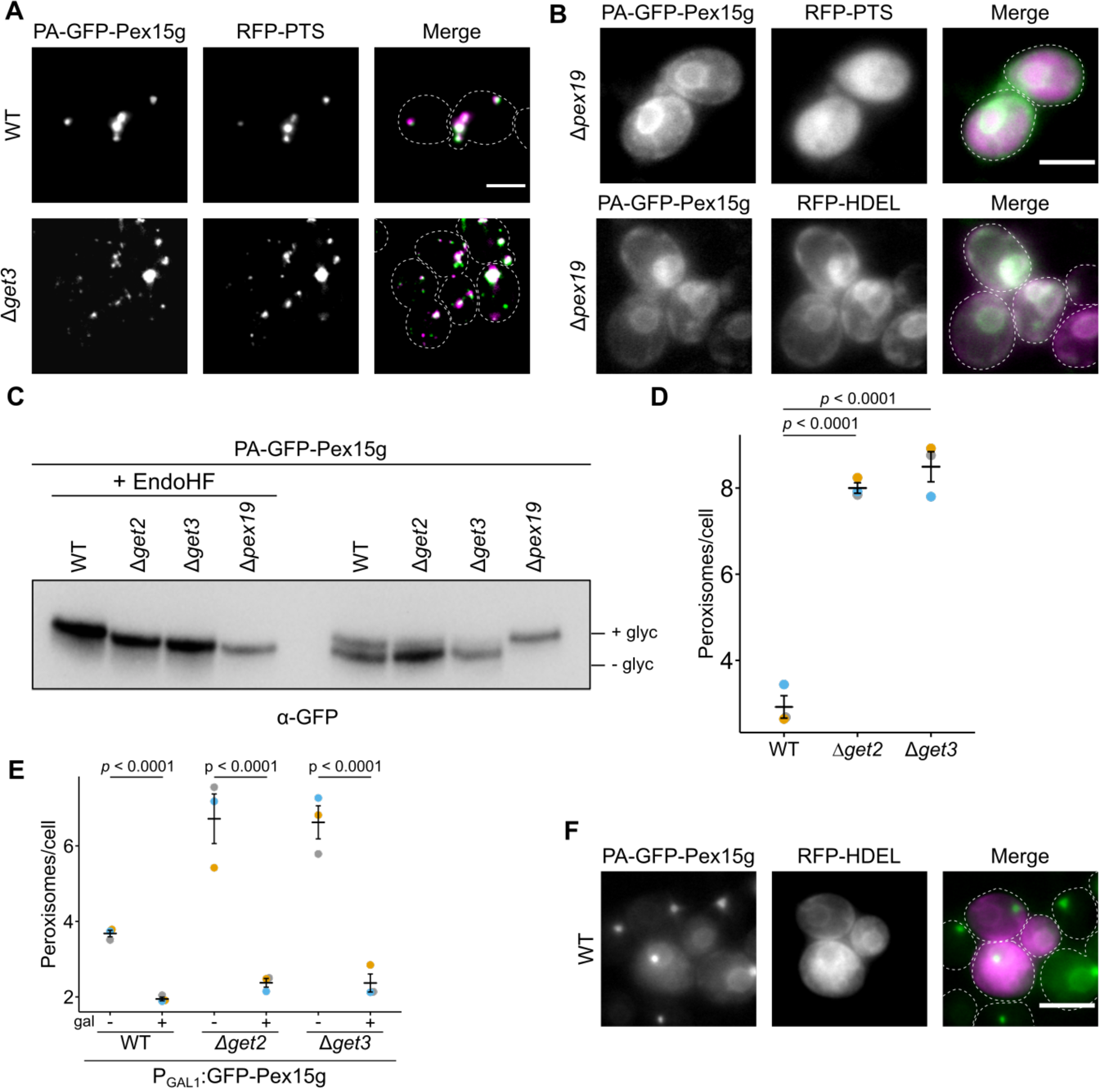
Dual targeting of Pex15 supports ER-peroxisome contacts. **(A)** Fluorescence microscopic images of wildtype (WT) and Δ*get3* cells expressing PA-GFP-Pex15g (green) and RFP-PTS1 (magenta). **(B)** Localization of PA-GFP-Pex15g (green) in in the ER in cells lacking the chaperone and targeting factor Pex19. RFP-HDEL (magenta) is a marker protein for the ER, RFP-PTS1 (magenta) is a marker for peroxisomes. Scale bar represents 5 µm **(C)** Proteins extracted from indicated strains were subjected to high resolution SDS-PAGE and immunoblot to visualize glycosylation of PA-GFP-Pex15g. **(D)** Quantification of peroxisome number of indicated strains expressing the peroxisomal marker mCherry-PTS1. Quantifications are based on *n* = 3 experiments. Each color represents one experiment. Error bars represent standard error of the mean. *P*-values were calculated using a two-sided unpaired Student’s *t*-test. **(E)** Quantification of peroxisome number of indicated strains either grown in glucose medium or galactose medium for 3 h to induce expression of GFP-Pex15g under control of a galactose inducible promoter. **(F)** Fluorescence microscopic images of cells co-expressing RFP-HDEL (magenta) PA-GFP-Pex15g (green) under control of the *MET25* promoter grown in the presence of 50 µM methionine. Scale bars represent 5 µm. Quantifications are based on *n* = 3 experiments. Each color represents one experiment. Error bars represent standard error of the mean. *P*-values were calculated using a two-sided unpaired Student’s *t*-test. Thus, trafficking of a distinct dual affinity protein may happen at an organelle contact site and promote this connection.

In wildtype cells expressing PA-GFP-Pex15g we found glycosylated and non-glycosylated forms of the chimeric protein, which together with the subcellular localization indicated dual targeting and transit (Fig. 6A and 6C). Deletion of genes coding for components of the GET complex led to a reduction of glycosylation and to the accumulation of small aberrant peroxisomes resembling those of Δ*pex5* and Δ*mdm10* cells (Fig. 6C; compare Fig. 6D to Figs. 1 and 5). Peroxisomal localization of PA-GFP-Pex15g was still observed (Fig. 6A) indicating that the phenotype does not result from lack of Pex15 on peroxisomes.

To test whether it is the ER transit of Pex15 that is required for maintaining regular peroxisomes, we overexpressed GFP-Pex15g via a galactose inducible promoter and observed suppression of the Δ*get2* and Δ*get3* phenotypes (Fig. 6E; Fig. S7) suggesting that mistargeting of Pex15 contributes to the generation of aberrant peroxisomes in mutants lacking a functional GET complex. Prolonged overexpression of PA-GFP-Pex15g resulted in peroxisomes attached to the ER (Fig. 6F) resembling peroxisomes attached to mitochondria by overexpression of dual affinity PTS1 proteins (Fig. 1).

### Synthetic genetics interactions reveal redundant modules for tethering via dual targeting at mitochondria and the ER

If tethering of peroxisomes to the ER can be supported by proteins sorted via the ER then this is a parallel or redundant way to create ER-peroxisome contacts.

Pex30 and orthologous proteins from other fungi are known to be involved in ER–peroxisome tethering, regulate membrane flux to peroxisomes, and act as a hub for peroxisome biogenesis at the ER [18,19, 45–49]. It was previously shown that there is a negative genetic interaction between *Δget2* and *Δpex30* [50, 51]. Similar to Δ*get2* and Δ*mdm10,* Δ*pex30* cells are characterized by an elevated number of smaller, highly mobile peroxisomes (Movie2) [45]. We observed substantial cytosolic mistargeting of mCherry-PTS1, reduced cell growth and altered mitochondrial morphology in Δ*get2*Δ*pex30* mutants (Figs. 7A and S8). Cells with intact mitochondria showed the lowest number of peroxisomes, however peroxisomes that remained frequently in proximity to mitochondria (Figs 7A and S8). We conclude that the formation of functional peroxisomes requires tethering facilitated by parallel mechanisms - Pex30 on the one hand and protein import via the GET complex on the other. Mitochondrial dysfunction in the double mutant may be caused by mistargeting of proteins designated for other organelles including peroxisomes.

**Figure 7.**
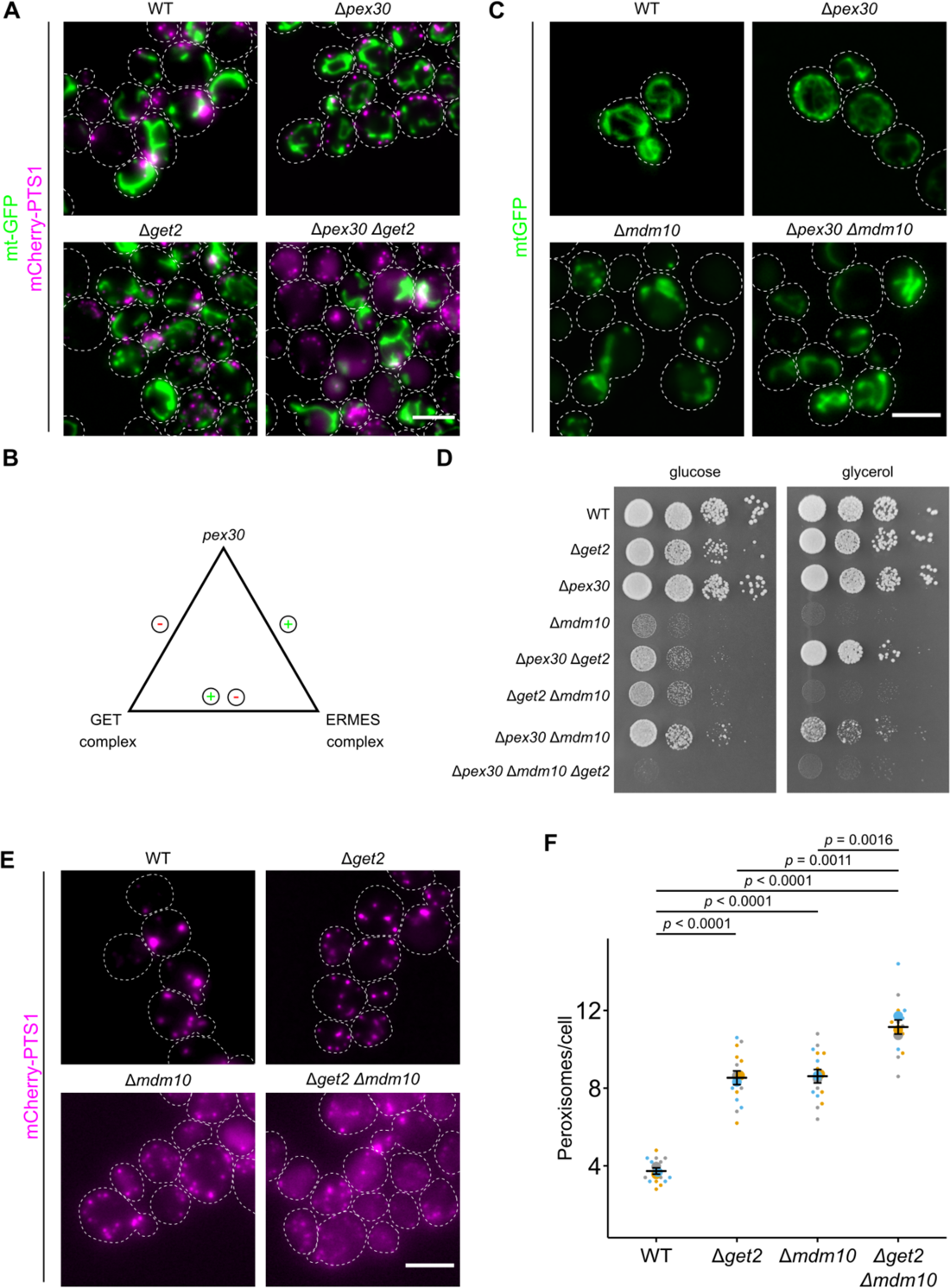
Genetic interactions between genes encoding tethering/targeting factors at mitochondria and the ER. **(A)** Fluorescence microscopic images of indicated strains expressing the peroxisomal marker mCherry-PTS1 and the mitochondrial marker mt-GFP. **(B)** Schematic illustration of the genetic interactions between *PEX30* and genes encoding components of ERMES and the GET complex according to Costanzo et al., 2010, 2016. Plus sign indicates a positive genetic interaction, minus signs indicate negative genetic interactions. **(C)** Fluorescence microscopic images of indicated strains expressing the mitochondrial marker mt-GFP. **(D)** Serial dilutions of logarithmically growing cells were spotted on indicated media and incubated at 30°C for two d (glucose) and 5 d (glycerol). **(E)** Fluorescence microscopic images of indicated strains expressing the peroxisomal marker mCherry-PTS1. **(F)** Quantification of peroxisome number of indicated strains. Quantifications are based on *n* = 3 experiments. Each color represents one experiment. Error bars represent standard error of the mean. *P*-values were calculated using a two-sided unpaired Student’s *t*-test. Scale bars represent 5 µm.

Further synthetic genetic interactions were detected in comprehensive interaction maps [50, 51] e.g. Δ*mdm10*Δ*pex30* (positive genetic or buffering) and Δ*mdm12Δget2* (negative genetic or synthetic lethal) (Fig. 7B). The positive genetic interaction between *mdm10* and *pex30* hints to some level of epistasis. Indeed, in a Δ*mdm10*Δ*pex30* strain both the mitochondrial morphology phenotype and the growth defect of Δ*mdm10* single mutants were partially suppressed showing that absence of Pex30 counteracts absence of Mdm10 (Fig. 7C and 7D). Glucose medium improved the growth of the *Δmdm10Δget2* double mutant compared to the Δ*mdm10* single mutant which speaks for separate functions of Mdm10 and Mdm12 beyond their common role in ERMES complex formation (Fig. 7B - 7D). Recently, entry of mitochondrial cargo into the ER via the GET complex was described [52]. Elevated ER targeting of mitochondrial proteins could contribute to the growth defect of Δ*mdm10* single mutants on glucose-containing medium (Fig. 7D). In addition to enhanced cytosolic localization of mCherry-PTS1, Δ*get2*Δ*mdm10* cells contained an increased number of small peroxisomes compared to each of the single mutants (Fig. 7E and 7F). These data are consistent with the hypothesis that ERMES and GET complex exhibit a similar role – both are involved in regulating peroxisome growth and division by sorting of dual affinity cargo.

## Discussion

We have found that sorting of proteins with targeting signals for two distinct membranes involves sites of interorganellar contact and can concomitantly support their stabilization. This conclusion derives from an analysis of the phenotypic similarity of Δ*get2,* Δ*mdm10, Δpex5* and Δ*pex30* strains with respect to peroxisome number and appearance (Fig. 8A), the phenotypes of several double mutants and from the characterization of different types of dual affinity proteins (Fig. 8B).

**Figure 8.**
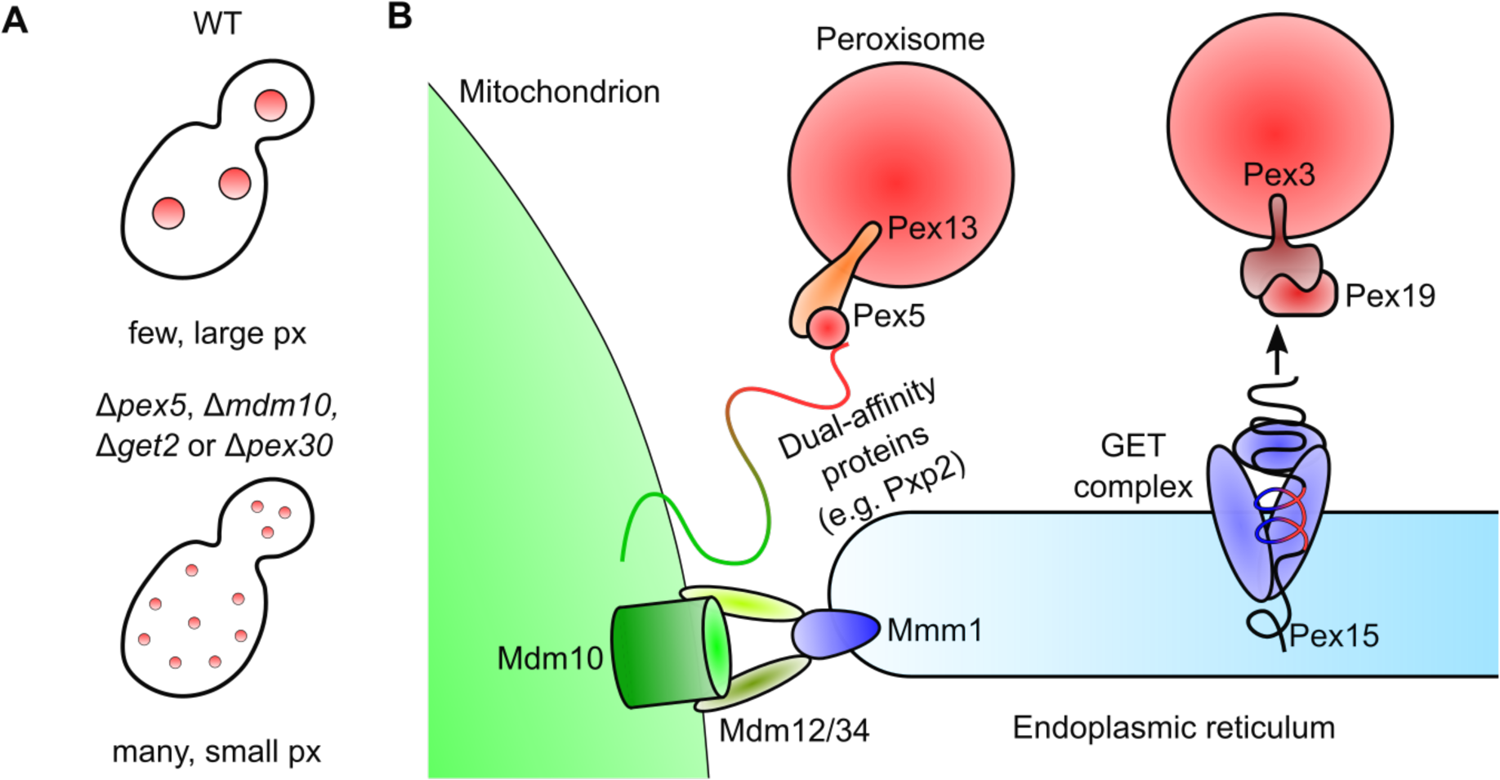
Model for sorting at contact sites and tethering via dual affinity proteins. **(A)** Deletion of genes encoding ERMES complex components (e.g. *MDM10*), GET complex subunits (e.g. *GET2*) *PEX5* or *PEX30* leads to the accumulation of smaller peroxisomes (px). **(B)** Model showing tethering of peroxisomes (red) to mitochondria (green) or the ER (blue). Formation of contacts to mitochondria may be achieved through simultaneous mitochondrial and peroxisomal targeting of dual affinity proteins (e.g. Pxp2). This correct sorting and tethering relies on the ERMES complex. Tethering of peroxisomes to the ER may occur through ER targeting of membrane proteins (e.g. Pex15 via the GET complex) and subsequent transit.

Such an interface may synchronize protein trafficking with organelle expansion, division and communication. Mobile targeting factors e.g. Pex5, Pex19 and Get3 and membrane complexes such as the ERMES and the GET complex are involved in this process. Depletion of a number of the targeting factors and membrane bound proteins directly or indirectly involved in the sorting caused accumulation of small peroxisomes (Fig. 8A). Overexpression of several dual affinity proteins led to a reduced number of peroxisomes and an increased fraction of peroxisomes attached to one or the other organelle. Although these proteins are a challenge for maintaining organelle identity [53, 54], they seem to be ideally suited to support organelle interactions by binding to different targeting factors and membrane bound translocation machineries or by trafficking from one organelle to the other. Dual affinity proteins may even define regions of reduced identity, which are likely to coincide with organelle contacts.

Localization of peroxisomal membrane proteins to mitochondria has been observed in mutants deficient in Pex19, components of the GET complex or the AAA-ATPase Msp1 [38,39,55]. Mitochondrial targeting of overexpressed Pex15 is observed upon deletion of components of the GET complex but not in wildtype cells [39], thus sorting via the ER probably represents a major avenue of transit. Other previous observations are consistent with our concept. Increased levels of Ant1, Fis1 and Pex11 enhanced PerMit contacts [16]. Ant1 and Pex11 accumulate in mitochondria of mutants lacking components required for peroxisomal membrane protein import [15, 17]. Another intriguing observation is that while many peroxisomal membrane proteins can target to peroxisomes without transitioning through the ER [56], several peroxisomal membrane proteins have evolved to be synthesized in vicinity to the ER and translocate from it [57]. Hence a potential reason for such rerouting may be for tethering purposes. One example is the integral membrane protein Pex3 that can either transit through the ER or directly insert into peroxisomes [58, 59]. Artificial targeting of Pex3 to mitochondria is compatible with peroxisome biogenesis in *S. cerevisiae* and provokes tethering of peroxisomes to this organelle [60]. Protein trafficking via a vesicular carrier [40,61–66] or direct membrane flow [24,25,67–70] accompanied by protein transit at contact sites may be parallel processes and may both depend on dual affinity proteins.

The decision to which organelle a peroxisome will preferentially attach can be influenced by the metabolic state of the cell [16]. In oleic acid medium proteins such as Tes1 and Cat2 are highly induced which may account for the observation of more frequent connections to mitochondria [16, 71].

Is this mode of targeting and tethering only true for peroxisomal proteins? The recently identified tether for mitochondria and the nuclear envelope Cnm1 shows hallmarks of a dual affinity protein comprising a targeting signal for the ER and for mitochondria at the N- and C-termini, respectively [72]. The discovery of an ER surface retrieval pathway for mitochondrial membrane proteins [73] exemplifies a related mechanism. The chaperone Djp1 critical in this pathway could function similar to Pex5 or Pex19 – the latter has unexpected role in sorting of mitochondrial TA proteins, as well [74]. Of interest, we detected Δ*djp1* among the mutants that decrease import of Ptc5-RFP-PTS to peroxisomes (Fig. 3B and 3C). It was shown previously that overexpression of Djp1 increases PerMit contacts [16] and that loss of Djp1 affects peroxisomal protein import [75]. The GET complex mediates import of peroxisomal and mitochondrial proteins into the ER [39, 52]. Several studies uncovered an overlap of factors for peroxisome and mitochondrial biogenesis in mammalian cells [76–78]. Accordingly, shared machinery controls protein targeting, biogenesis and division of both organelles and may contribute to tethering as well.

Upon discovery of ERMES as a tether structure, a link between protein transport, membrane flux and tethering mediated by the mitochondrial import protein Mdm10 was suggested [32]. Coupling of protein and membrane trafficking is a common principle in the secretory pathway [79] and it might also occur for peroxisomes at different contact sites. Simple predictions are difficult to make for redundant pathways based not on a single targeting factor but on the sum of interactions created by dual targeting of many proteins each with a unique affinity to the different targeting factors. In respect to the peroxisome, the functional redundancy of potential membrane sources and contact sites is an obstacle in classifying mutants defective in peroxisome biogenesis. A unifying hypothesis is complicated by the fact that several proteins can be directly imported into peroxisomes or traffic via a different organelle. We conclude that the dynamic interplay of membranes at interorganellar contact sites is influenced by targeting of proteins with dual localization signals.

## Acknowledgements

We thank Marisa Piscator for technical assistance in the Bölker laboratory. We recognize Bob Lesch for organizing data and Claudia Morales for support in the Schekman laboratory. We thank Amir Fadel and Lihi Gal for help with the automated library preparation and robotic screening procedures in the Schuldiner laboratory. We acknowledge Martin Thanbichler for sharing his microscope for time-lapse studies. We are grateful to Christof Taxis, Christian Renicke, Helle Ulrich, Jodi Nunnari, Michal Skruzny Ralf Erdmann and Roland Lill for antibodies, plasmids and strains. We thank Uwe Maier and Björn Sandrock for comments on the manuscript. TS received funding from a fellowship from DAAD. The project in the Schuldiner laboratory was supported by funding from the European Research Council (ERC) under the European Union’s Horizon 2020 research and innovation programme (Grant agreement No. 864068). The robotic system of the Schuldiner laboratory was purchased through the support of the Blythe Brenden-Mann Foundation. MS is an Incumbent of the Dr. Gilbert Omenn and Martha Darling Professorial Chair in Molecular Genetics. RS is an investigator of the Howard Hughes Medical Institute. JF was supported by a fellowship from Leopoldina and by the German research foundation (FR 3586/2-1).

## Methods

### Yeast strains, plasmids, and oligonucleotides

All *S. cerevisiae* strains used in this study have the genetic background of BY4741 or BY4742 [80] and are listed in supplementary table 4. Yeast cells were transformed as previously described using Li-acetate, PEG and ssDNA [81]. Gene replacement or endogenous tagging was carried out using PCR-amplification of cassettes based on an established toolbox system. The following plasmids were used for PCR amplification.: pKS133, pUG72, pHyg-AID*-6HA, pYM25, pCR125 and pYM-N22 [27,82–84]. Plasmids were generated by Gibson Assembly, recombinational cloning or classic molecular cloning [85–87]. Plasmids used or generated and oligonucleotides are listed in in supplementary table 4 [10,40,82,88–91].

### Yeast media and growth conditions

Strains were grown in either YPD (1% yeast extract, 2% peptone, 2% glucose) medium or synthetic medium containing 2% glucose, 0.67% yeast nitrogen base without amino acid, 0.5% ammonium sulfate and complementary supplement mixtures (CSM) or drop-out mixtures lacking appropriate amino acids when experiments required selection. For galactose induction of RFP-Pex15g, cells were grown overnight in synthetic medium containing 2% raffinose, 0.1% glucose, 0.67% yeast nitrogen base without amino acids, 0.5% ammonium sulfate and CSM-His or CSM-His-Leu overnight. Cells were washed once with H_2_O and diluted in 2% raffinose, 0.67% yeast nitrogen base without amino acids and CSM-His or CSM-His-Leu. After 2h 2% galactose was added and cells were incubated for 3 h. For regulation of Pex5 expression, cells were grown overnight in YP (1% yeast extract, 2% peptone) medium containing 2% raffinose and 0.1% glucose. Cells were washed once with H_2_O and diluted to an OD_600_ of 1 in YP containing 2% raffinose. After 2 h, cells were harvested and resuspended in YP medium containing 2% galactose. After 45 min, cells were again harvested and resuspended in YPD medium. Samples were taken at indicated time points and analyzed either by fluorescence microscopy or SDS-PAGE and immunoblot. For depletion of Pex13, cells were precultured in either YPD or synthetic medium containing 2% glucose, 0.67% yeast nitrogen base without amino acids, 0.5% ammonium sulfate and dropout-mix lacking histidine. Cells were diluted in the appropriate medium to an OD_600_ of 0.5 and incubated at 30°C for 2 h. The culture was then split into two aliquots of equal volume and one culture was supplemented with 2 mM indole-3-acetic acid (auxin). Both cultures were incubated at 30°C and at indicated time points samples were taken and analyzed by either fluorescence microscopy or SDS-PAGE and immunoblot. For growth assays on plates 5 µl of serial tenfold dilutions of logarithmically growing cells (starting OD_600_ = 1) were spotted onto solid synthetic media either containing 2% glucose or 2% glycerol and were incubated for 2 – 5 days at 30°C. Cells expressing PA-GFP-Pex15g under control of a methionine regulatable promotor (Lam et al., 2010) were grown in the presence of 200 µM methionine or 50 µM methionine to achieve substantial overexpression.

### High content screening

To create collections of haploid strains, we constructed a Synthetic Genetic Array (SGA) compatible query strain co-expressing the peroxisomal marker Pex3-GFP (endogenous) together with Ptc5-RFP-PTS1 (overexpression). Using automated mating approaches the query strain was crossed with deletion and DAmP arrayed collections [29, 30]. Yeast manipulations in high-density format were performed on a RoToR bench top colony arrayer (Singer Instruments). In short: mating was performed on rich medium plates, selection for diploid cells was performed on SD monosodium glutamate (MSG) plates containing geneticin (Formedium, 500 µg ml^−1^), nourseothricin (WERNER BioAgents “ClonNat”, 200 µg ml^−1^) and hygromycin B (Formedium, 500 µg ml^−1^). Sporulation was induced by transferring cells to nitrogen starvation media plates for 7 d. Haploid cells containing the desired mutations were selected by transferring cells to SD MSG plates containing geneticin, neurseothricin and hygromycin B, alongside the toxic amino-acid derivatives canavanine and thialysine (Sigma-Aldrich) to select against remaining diploids, and lacking leucine to select for spores with an alpha mating type. The collections were visualized using an automated microscopy setup. In brief, cells were transferred from agar plates into 384-well polystyrene plates for growth in liquid media using the RoToR arrayer. Liquid cultures were grown in a LiCONiC incubator overnight at 30 °C in SD medium (6.7 g l^−1^ yeast nitrogen base with ammonium sulfate and 2% glucose) supplemented with complete amino acids. A JANUS liquid handler (PerkinElmer) connected to the incubator was used to dilute the strains to an OD_600_ of ∼0.2. Plates were incubated at 30 °C for 4 h in SD medium. The cultures in the plates were then transferred by the liquid handler into glass-bottom 384-well microscope plates (Matrical Bioscience) coated with concanavalin A (Sigma-Aldrich). After 20 min, wells were washed twice with SD-Riboflavin complete medium to remove non-adherent cells and to obtain a cell monolayer. The plates were then transferred to a ScanR automated inverted fluorescent microscope system (Olympus) using a robotic swap arm (Hamilton). Images of cells in the 384-well plates were recorded in SD-riboflavin at 24 °C using a ×60 air lens (NA 0.9) with an ORCA-ER charge-coupled device camera (Hamamatsu). Images were acquired in two channels: GFP (excitation filter 490/20 nm, emission filter 535/50 nm) and mCherry (excitation filter 572/35 nm, emission filter 632/60 nm). After acquisition, images were manually reviewed using the ScanR analysis program and ImageJ.

### Preparation of post-nuclear supernatants and differential centrifugation

For preparation of crude organelles, 3-6 l cultures were grown in synthetic medium containing appropriate amino acids for selection to logarithmic phase (OD_600_ = 0.8-1) at 30°C. Cells were harvested and suspended in 25 ml 100mM Tris/HCl buffer (pH 9.4) per liter of starting culture containing 10 mM DTT. The suspension was incubated for 10 min at room temperature and centrifuged at 600 x *g* for 5 min. Cell pellets were suspended in 25 ml lyticase buffer (0.7M sorbitol, 0.75xYP, 0.5% glucose, 10mM Hepes/OH, 1mM DTT; pH 7.4) per liter of starting culture and lyticase (10^5^ units per liter starting culture) was added. Suspensions were incubated for 30 – 45 min at 30°C and the efficiency of spheroplast formation was determined by measuring the decline of OD_600_ after suspension of samples in H_2_O. Spheroplasts were washed 3 times with 25 ml 2xJR buffer (0.4M sorbitol, 100mM KOAc, 40mM Hepes/OH (pH 7.4), 4mM EDTA, 2mM DTT) per liter starting culture, suspended in 5 ml 2xJR (containing a protease inhibitor cocktail: 1mM 4-aminobenzamidinedihydrochloride, 1 µg/ml aprotinin, 1µg/ml leupeptin, 1mM phenylmethylsulfonyl fluoride, 10 µg/ml N-tosyl-L-phenylalanine chloromethyl ketone and 1 µg/ml pepstatin) per liter starting culture and frozen at −80°C. Cells were thawed in iced water and disrupted by 20 strokes with a Potter-Elvehjem homogenizer. Nuclei were removed from homogenates by centrifugation two times at 600 x *g*. For differential centrifugation, 1 mg PNS fraction was used. PNS (100 µl) fractions were centrifuged at 13k x g for 5 min. The resulting supernatant was centrifuged at 100k x g for 20 min. All fractions were analyzed by SDS-PAGE and immunoblot in amounts representing the initial volume.

### Density gradient centrifugation

Post-nuclear supernatants (PNS) for density gradient centrifugation were prepared using a modified protocol from Cramer et al., 2015 (steps 1–16). Cells were precultured in 500 ml YP medium supplemented with 0.1% glucose at 30 °C overnight. Cells were harvested at 6000 × g for 6 min, washed once with 30 ml sterile water, and resuspended in 500 ml YNBO medium. After incubation at 30 °C for 16 h, cells were harvested as described above, washed twice with 30 ml sterile water, and incubated in 15 ml DTT buffer (100 mM Tris, 10 mM DTT, pH 7.4) for 30 min at 30 °C with gentle agitation. Cells were sedimented by centrifugation at 600 × g, washed three times with 15 ml sorbitol buffer (20 mM HEPES, 1.2 M sorbitol), and incubated with 15 ml sorbitol buffer containing 20 mg Zymolyase 100 T (Roth) at 30 °C with gentle agitation. Digestion of yeast cell walls was monitored photometrically at OD600 and digestion was stopped when at least half of the cells lysed upon addition of water. From this point all steps were carried out on ice or at 4 °C. Spheroplasts were gently washed three times with 15 ml sorbitol buffer, resuspended in 15 ml lysis buffer (5 mM MES, 0.5 mM EDTA, 1 mM KCl, 0.6 M sorbitol, 1 mM 4-aminobenzamidine-dihydrochloride, 1 µg/ml aprotinin, 1 µg/ml leupeptin, 1 mM phenylmethylsulfonyl fluoride, 10 µg/ml N-tosyl-L-phenylalanine chloromethyl ketone, and 1 µg/ml pepstatin), and frozen at −80 °C overnight. Spheroplasts were thawed on ice and homogenized using a Potter-Elvehjem homogenizer (2 × 12 strokes). Nuclei and cell debris were removed by two subsequent centrifugations at 1600 × g for 10 min. Subsequently, the PNS was diluted to an OD600 of 1, aliquoted and frozen at −80 °C. A total of 200 µl of PNS without cytosol were loaded onto a NycoDenz density gradient consisting of 333 µl 20%, 666 µl 25%, 666 µl 30%, and 333 µl 35% NycoDenz in gradient buffer A (5 mM MES, 1 mM EDTA, 1 mM KCl, and 0.1% (v/v) ethanol). Gradients were centrifuged in a Beckman L7-65 ultracentrifuge equipped with a Sorvall TST 60.4 rotor at 100,000 × g (35,000 rpm) for 90 min at 4 °C. Twelve 183 µl fractions were collected from the top of the gradient and used for SDS–PAGE and immunoblot.

### Epifluorescence microscopy

A total of 200 µl of hot 1.5% agarose melted in water was used to create a thin agarose cushion on a 76 × 26 mm microscope slide (Roth, Karlsruhe, Germany). Cells were washed with water, concentrated 10-fold, and 3 µl aliquots were spotted onto the middle of the agarose pad and covered with an 18 × 18 mm coverslip (Roth, Karlsruhe, Germany). Microscopy was performed on an Axiovert 200 M inverse microscope (Zeiss) equipped with a 1394 ORCA ERA CCD camera (Hamamatsu Photonics), filter sets for cyanGFP, enhanced GFP (EGFP), yellow fluorescent protein (YFP), and rhodamine (Chroma Technology, Bellows Falls, VT), and a Zeiss 63× Plan Apochromat oil lens (NA 1.4). Single-plane bright field or phase contrast images and *z*-stacks of the cells (0.5 µm *z*-spacing) in the appropriate fluorescence channels were recorded, using the image acquisition software Volocity 5.3 (Perkin-Elmer). Images were processed and evaluated in ImageJ [92]. Alternatively, images were taken on an inverted Zeiss Axioobserver fluorescence microscope and analyzed with Metamorph (Molecular Devices). For protein localization analysis, *z*-projections of deconvolved image stacks of the fluorescent channels were used. Deconvolution was performed on the *z*-stacks by the ImageJ plugin DeconvolutionLab with 25 iterations of the Richardson–Lucy algorithm [93]. Pearson’s correlation coefficients were determined with Volocity 5.3.

### Automated time lapse microscopy

Exponentially growing liquid cultures were spotted on 1.5 % agarose pads containing synthetic media. Images were taken with a Zeiss Axio Observer. Z1 microscope equipped with a ×100/1.46 Oil DIC objective and a pco.edge 4.2 sCMOS camera (PCO). An X-Cite 120PC metal halide light source (EXFO, Canada) and ET-YFP or ET-TexasRed filter cubes (Chroma, USA) were used for fluorescence detection. Images were recorded with VisiView 3.3.0.6 (Visitron Systems) and processed with Metamorph 7.7.5 (Molecular Devices) and Adobe Illustrator CS6 (Adobe Systems). Images were taken every hour and cells were incubated at 30°C.

### Immunoblotting and antibodies

Small amounts of protein were extracted as described by Kushnirov (2000) using a modified sample buffer (50 mM Tris-HCl, pH 6.8, 2% SDS, 6% glycerol, 0.025% bromophenol blue, and 50 mM dithiothreitol) [94]. In brief, 1 OD_600_ of yeast cells were centrifuged at 13,000 × *g* for 1 min and incubated with 300 µl 0.2 M NaOH for 5 min at room temperature. Cells were centrifuged, resuspended in 50 µl sample buffer, incubated for 5 min at 95 °C, centrifuged again, and the supernatant was transferred to a new reaction tube. Proteins were incubated for 5 min at 95 °C and rotated at 750 rpm prior to loading. SDS–PAGE was performed with self-cast or Midi-protean TGX precast gels (BioRad), PageRuler Prestained protein ladder (ThermoFisher) as protein standard and a BioRad Mini- or Midi-Protean cell. Proteins were blotted on PVDF membranes in a BioRad Mini Trans-Blot cell or Criterion Blotter at 30 V for 16 h at 4°C or at 70 V for 1h at 4°C. Membranes were blocked for 30 min in TBST containing 5% nonfat dry milk and incubated in primary antibody containing 0.02% sodium azide with gentle agitation for 2 h at room temperature or overnight at 4 °C. After removal of the antibody solution, membranes were washed five times with TBST for 5 min, incubated with HRP-conjugated secondary antibody for 45 min at room temperature, washed five times with TBST for 5 min, and then developed using either Pierce ECL Immunoblotting Substrate (ThermoFisher) or SuperSignal West Pico PLUS Chemiluminescent Substrate (Thermo Fisher). The following antibodies were used in this study: anti-GFP (1:2000; TP401, Torrey Pines Biolabs), anti-HA (1:2000; ab1302275, Abcam), anti-tagRFP (1:1000; AB233, Evrogen), anti-Por1 serum (1:2000; kindly provided by Roland Lill, Marburg), anti-Pex5 serum (1:10.000; kindly provided by Ralf Erdmann, Bochum), anti-Kar2 serum (1:5000), anti-Pgk1 (1:2000; 22C5D8, ThermoFisher), anti-mCherry (1:1000; TA150125, ThermoFisher), anti-Protein A (1:2000; ab19483, Abcam), goat anti-mouse IgG-HRP (1:10.000 - 1:50.000; 31430, ThermoFisher), goat anti-rabbit IgG-HRP (1:10.000 – 1:50.000; 31460, ThermoFisher).

### EndoHF treatment

Protein extract samples (5µl) were subjected to EndoHf (New England Biolabs) treatment according to manufactures instructions. EndoHf (4µl, 4000 units) was used per reaction. To resolve different mobility of glycosylated and non-glycosylated proteins, we used 7.5% Tris-glycine midi-gels (Bio-Rad).

### Statistics and reproducibility

Microscopic data was collected from 3 independent *S. cerevisiae* cultures. Five images per culture were quantified. Pearson’s correlation coefficient was calculated with Volocity 5.5.2. Superplots [95] and Student’s *t*-tests were computed using RStudio 1.2.1335 with R 3.6.0. Blots are structured as follows: center line, mean; error bars, standard error of the mean; big circles, mean of experiments; and small circles, data points of experiments. *P*-values were calculated using an unpaired, two-sided Student’s *t*-test. For calculation of Pearson’s correlation coefficients all analyzed images contained ten or more cells and one image represents one data point. For quantification of contacts between mitochondria and peroxisomes five images per strain with at least five cells from 3 independent experiments were analyzed. Quantification was performed by manual inspection of cells. Peroxisomes were counted as mitochondria associated if no black pixels were detected between fluorescent signals of respective marker proteins. Inspection was carried out without knowledge of the genotypes. All experiments were at least repeated three times with similar results.

---

**Figure S1.**
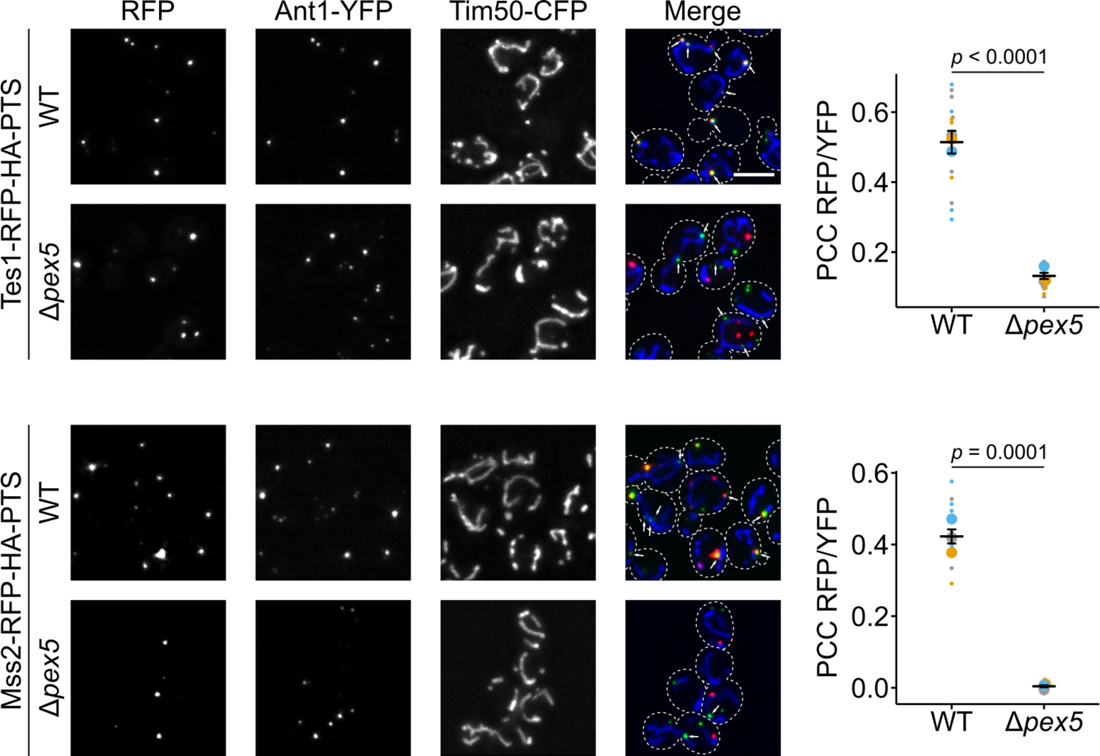
Peroxisomal localization of Tes1 and Mss2 depends on *pex5*. Tes1-RFP-HA-PTS (red) or Mss2-RFP-HA-PTS (red) were co-expressed with the peroxisomal membrane protein Ant1-YFP (green) and mitochondrial inner membrane protein Tim50-CFP (blue) in indicated strains (left). Subcellular localization was determined by fluorescence microscopy. Scale bar represents 5 µm. White arrows indicate peroxisomes proximal to mitochondria. Quantification of correlation between Tes1-RFP-HA-PTS (top right) signal or Mss2-RFP-HA-PTS (bottom right) signal and Ant1-YFP signal in indicated strains. PCC refers to Pearson’s correlation coefficient. Quantifications are based on *n* = 3 experiments. Each color represents one experiment. Error bars represent standard error of the mean. *P*-values were calculated using a two-sided unpaired Student’s *t*-test.

**Figure S2.**
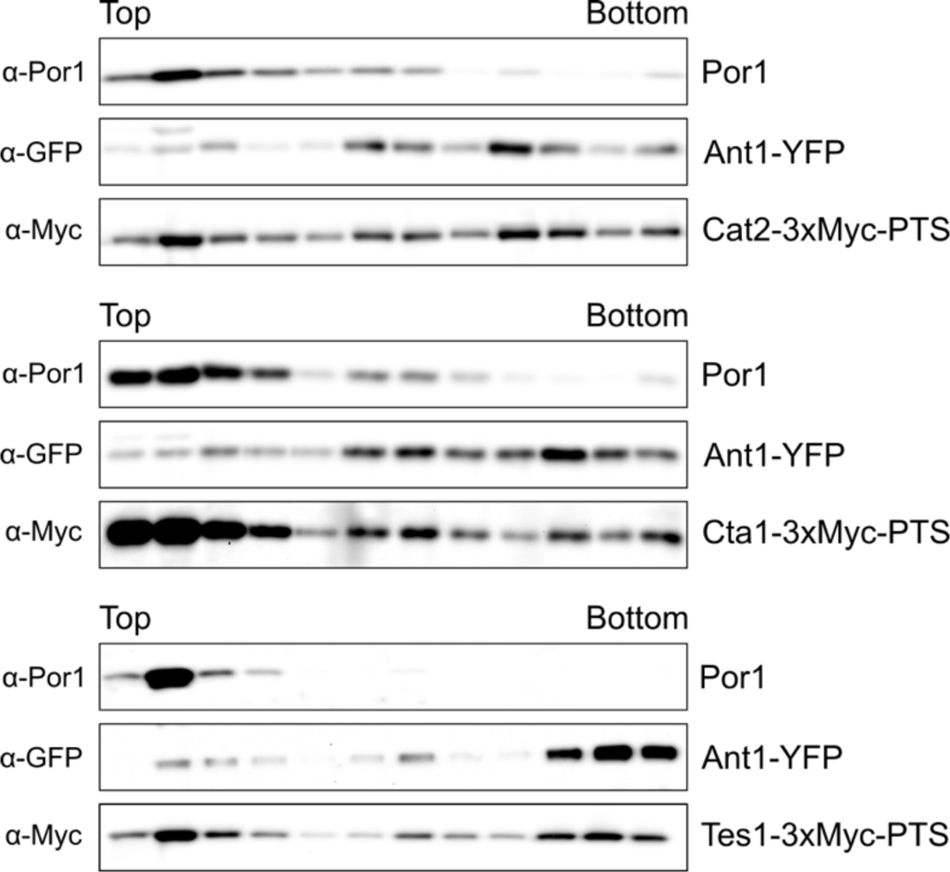
Subcellular localization of endogenously tagged variants of dually targeted PTS1 proteins in wildtype cells expressing Ant1-YFP and Tim50-CFP. Subcellular localization of endogenously tagged variants of dual affinity proteins was determined by density gradient centrifugation. Twelve fractions, collected from the top of the gradient, were analyzed by SDS–PAGE and Western blot. Ant1-YFP is a peroxisomal membrane protein and Por1 is localized in the mitochondrial outer membrane. **Movie 1**: **Movement of peroxisomes in indicated strains.** Frames were captured with the following excitation times and gain settings: excitation time, 400ms; gain, 228

**Fig. S3.**
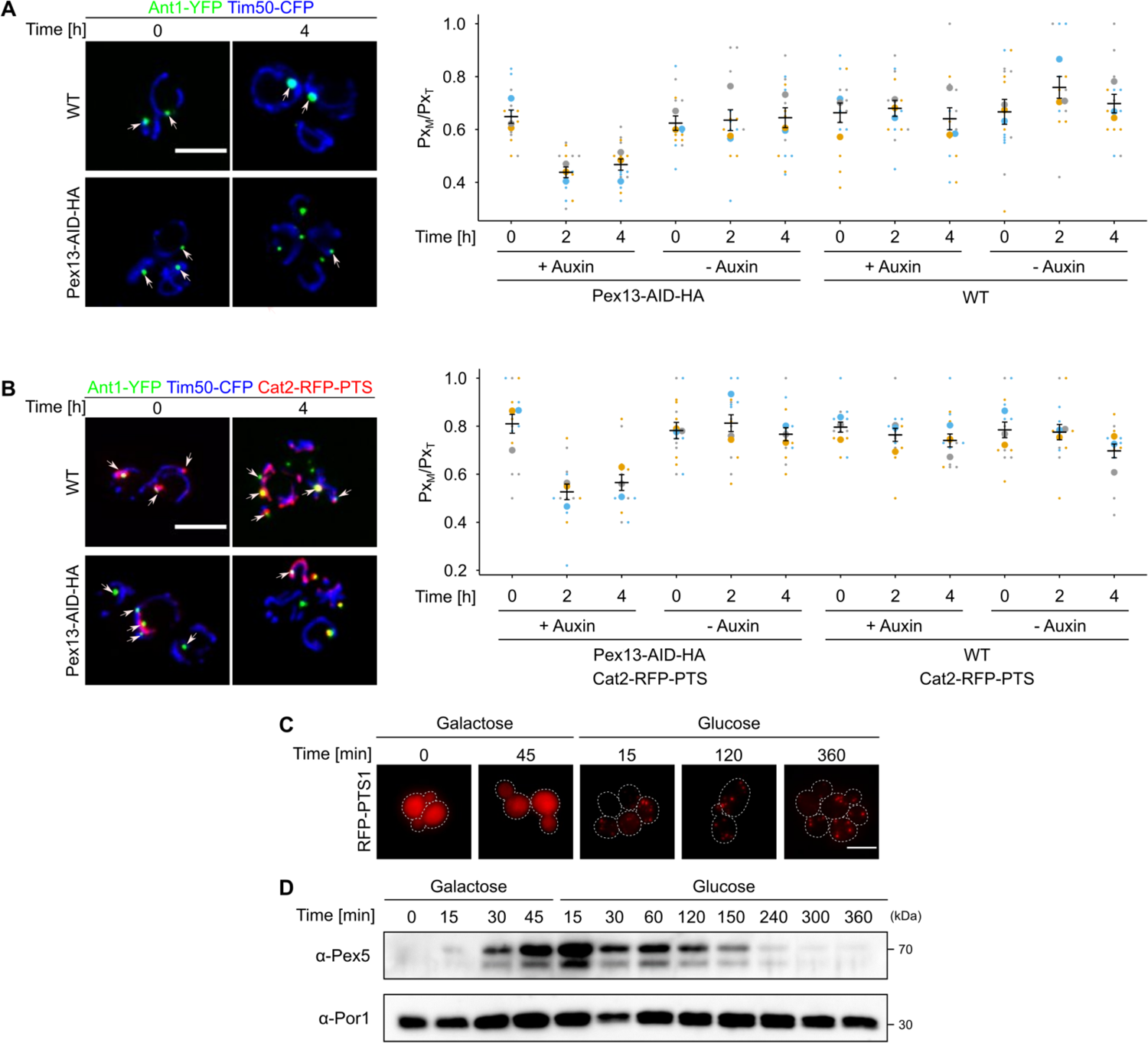
Decrease of PerMit contacts via conditional mutants. **(A)** Fluorescence microscopic images of indicated strains expressing Ant1-YFP (green) and Tim50-CFP (blue) in the presence of 2 mM indole-3-acetic acid at indicated time points (left). Arrows indicate peroxisomes that contact mitochondria. Quantification of the fraction of peroxisomes in contact with mitochondria (Px_M_) relative to the total peroxisome count (Px_T_) of the indicated strain at the indicated time points and growth conditions (right). Quantifications are based on *n* = 3 experiments. Each color represents one experiment. Error bars represent standard error of the mean. **(B)** Identical to (A), except that the cells also expressed Cat2-RFP-PTS1 (red) leading to increased initial contacts prior to depletion. **(C)** Fluorescence microscopic images of a galactose chase and shut-off experiment using a galactose-inducible conditional *pex5* mutant expressing RFP-PTS1. **(D)** Pex5 levels from the indicated growth conditions at indicated time points of the strain described in figure 2I were analyzed by SDS-PAGE and Immunoblot. Por1 served as loading control. Scale bars represents 5 µm.

**Figure S4.**
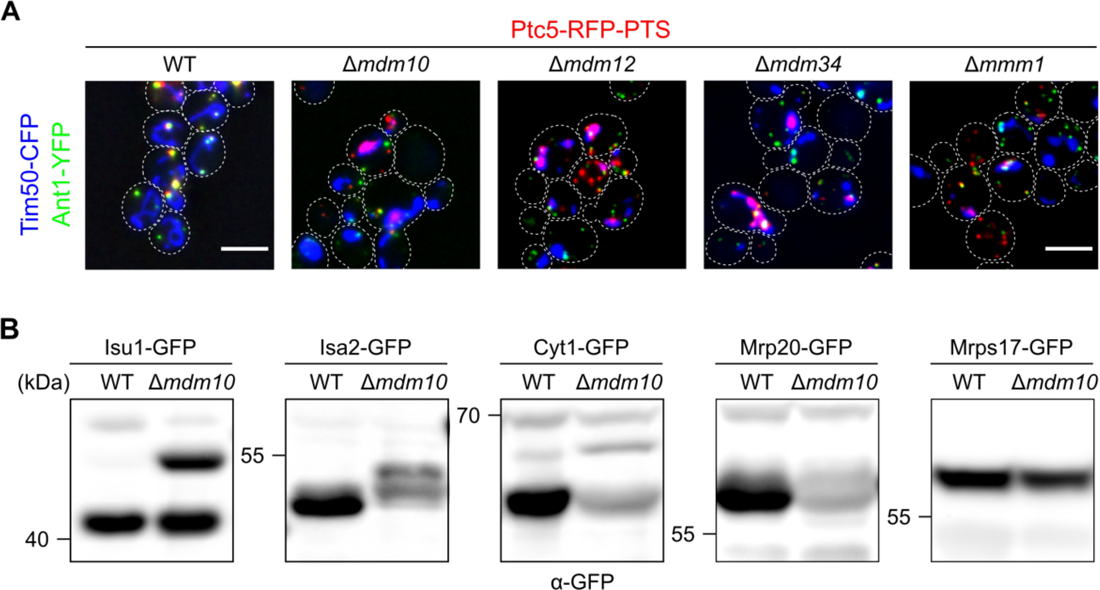
Lack of ERMES components affects mitochondrial and peroxisomal protein import. **(A)** Subcellular localization of Ptc5-RFP-PTS (red), the peroxisomal membrane protein Ant1-YFP (green) and the mitochondrial inner membrane protein Tim50-CFP (blue) in indicated strains was analyzed using fluorescence microscopy. Scale bar represents 5 µm. **(B)** *Bona fide* mitochondrial proteins C-terminally tagged with GFP at the endogenous locus in wildtype (WT) and Δ*mdm10* cells were analyzed. Whole cell extracts were subjected to SDS-PAGE followed by immunoblot. Concentrations of protein extracts were adapted to each other to focus on processing.

**Figure S5.**
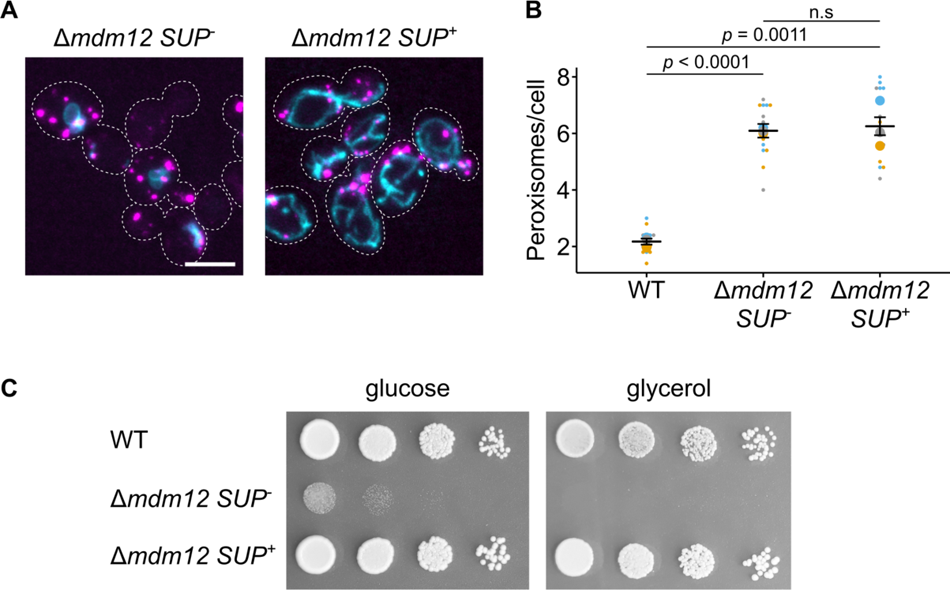
Suppression of Δ*mdm12* does not suppress formation of aberrant peroxisomes. **(A)** Fluorescence microscopic images of Δ*mdm10* cells (*SUP^-^*) or Δ*mdm10* cells with a suppressor mutation (*SUP^+^*) co-expressing the peroxisomal marker Ant1-YFP (magenta) and the mitochondrial inner membrane protein Tim50-CFP (cyan). Scale bar represents 5 µm. **(B)** The number of peroxisomes per cell was quantified in the indicated strains. Quantifications are based on *n* = 3 experiments. Each color represents one experiment. Error bars represent standard error of the mean. *P*-values were calculated using a two-sided unpaired Student’s *t*-test. **(C)** Serial dilutions of logarithmically growing cells of indicated strains were spotted on indicated media and incubated at 30°C.

**Figure S6.**
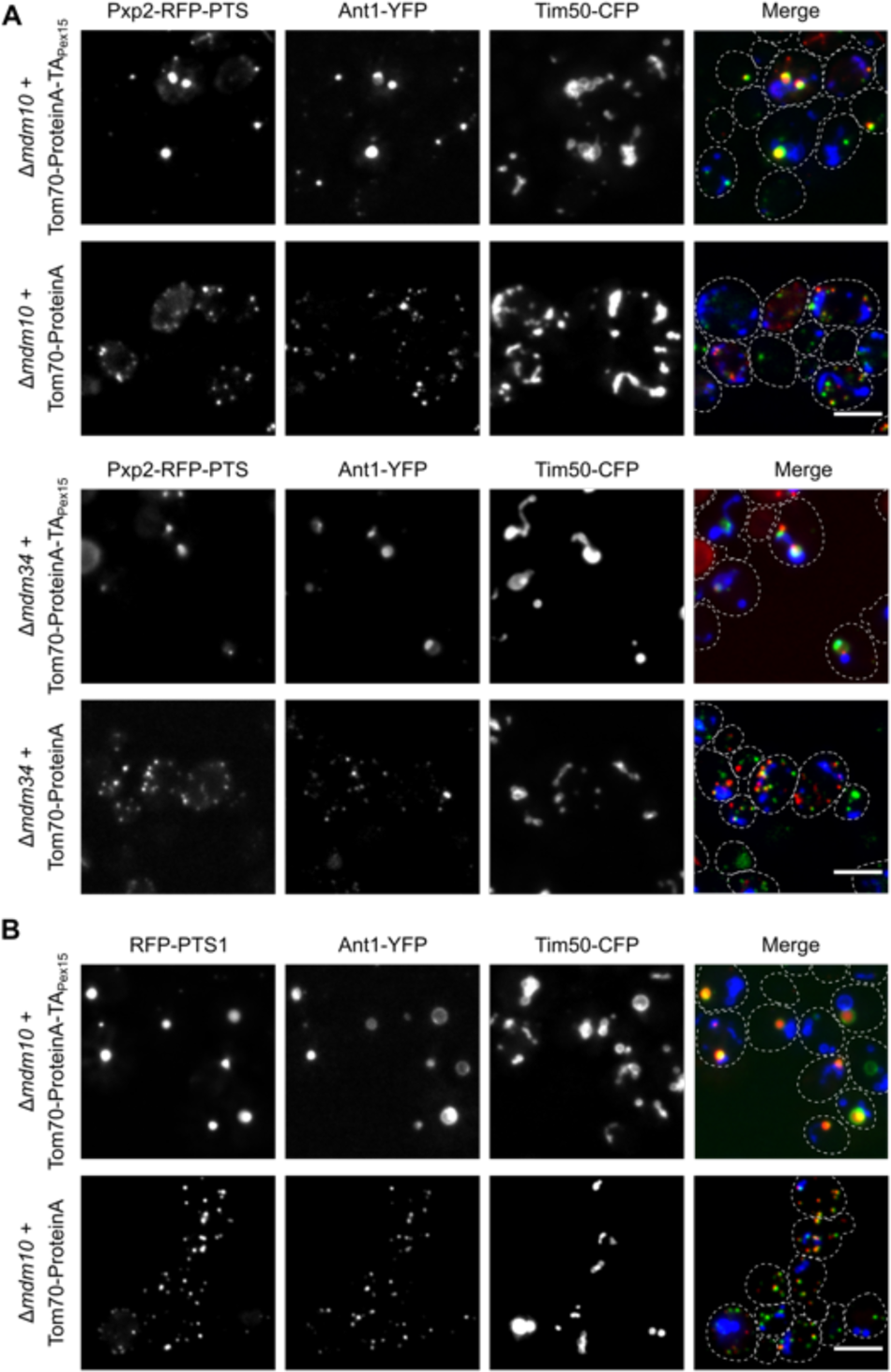
Analysis of peroxisomes in ERMES mutants upon expression of a synthetic tether. Strains deleted for *mdm10* or *mdm34* co-expressing Pxp2-RFP-PTS1. **(A,** red) RFP-PTS1 **(B,** red) or the peroxisomal membrane protein Ant1-YFP (green) and the mitochondrial protein Tim50 (blue) either in the presence of Tom70-ProteinA-TA_Pex15_ or a control protein. Scale bars represent 5 µm.

**Figure S7.**
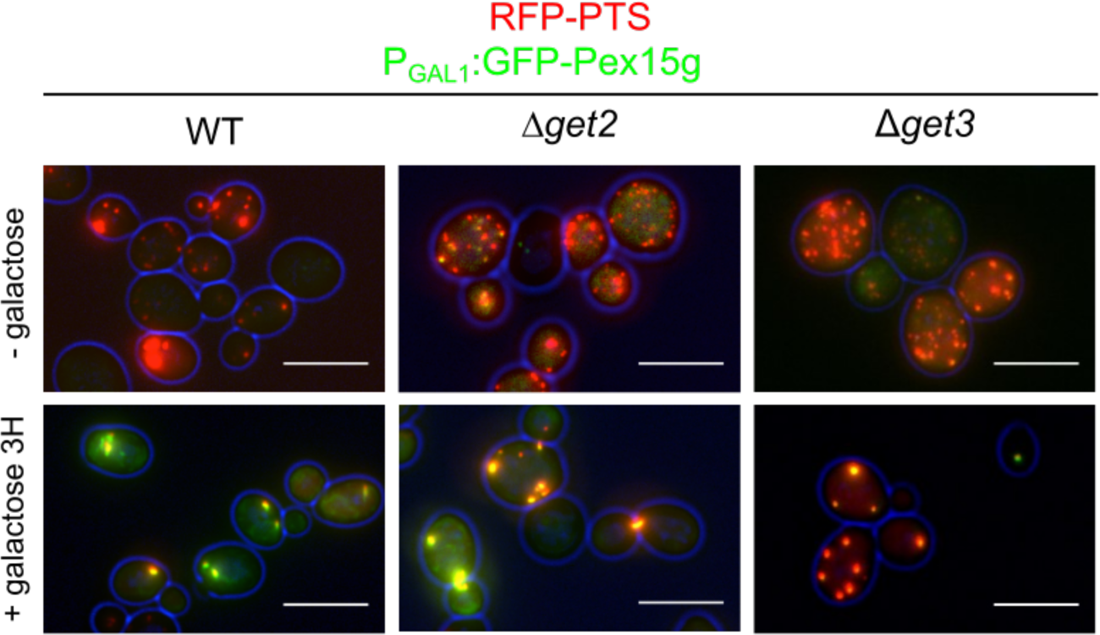
Suppression of the *get* deletion phenotypes by overexpression of GFP-Pex15g. Fluorescence microscopic images of cells co-expressing GFP-Pex15g (green) under control of a galactose inducible promoter and RFP-PTS1 (red). Scale bars represent 10 µm. Upper panel: glucose grown cells; lower panel: 3 h galactose induction. **Movie 2. Movement of peroxisomes in indicated strains.** Frames were captured with the following excitation times and gain settings. WT, 75 ms, gain 23; Δ*pex30*, 80 ms, gain 79; Δ*get2*, 100 ms, gain 63; Δ*mdm10*; 100 ms, gain 48.

**Figure S8.**
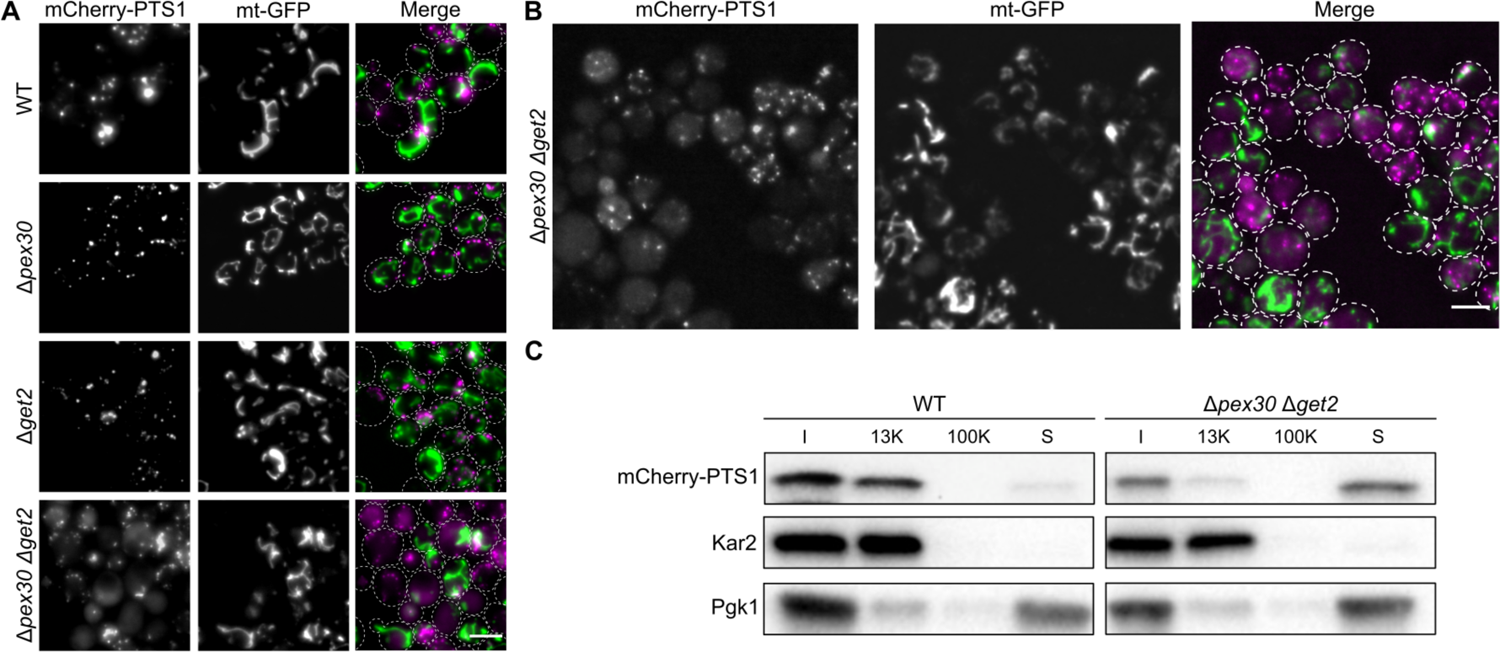
Depletion of *PEX30 and GET2* affects peroxisomes and mitochondria. **(A and B)** Fluorescence microscopic images of strains expressing the mitochondrial marker mt-GFP (green) and the peroxisome marker mCherry-PTS1 (magenta) in indicated mutants. Scale bar represents 5 µm. **(C)** Differential centrifugation of post nuclear supernatants of indicated strains. Fractions were analyzed by SDS-PAGE and immunoblot. I, input; 13k, 13k x *g* pellet; 100k, 100k *x* g pellet; S, 100k x *g* supernatant. Pgk1 is a cytosolic protein, Kar2 an ER resident protein.

